# Plectin Regulates Focal Adhesion Dynamics and Cytoskeletal Organization in Mouse Astrocytes: Implications for Reactive Astrogliosis

**DOI:** 10.1101/2025.01.14.632892

**Authors:** Borut Furlani, Maja Potokar, Victorio Martin Pozo Devoto, Gerhard Wiche, Robert Zorec, Jernej Jorgačevski

## Abstract

Reactive astrogliosis, a hallmark of central nervous system pathologies, involves cellular responses, including morphological remodelling and upregulation of intermediate filaments such as vimentin. These changes are driven by cytoskeletal dynamics and are mediated by focal adhesions (FAs). Our study identifies plectin, a versatile cytoskeletal linker protein, as a critical modulator of FA-associated processes in mouse astrocytes. We demonstrate that plectin localizes to astrocyte FAs, where it regulates their number, maturation, turnover, and the mobility of FA components. Plectin also polarizes within FAs, depending on their maturation state, and controls the recruitment of key cytoskeletal elements particularly vimentin. In plectin-deficient astrocytes, the vimentin network exhibits impaired connectivity, accompanied by altered viscoelastic properties of the cells. In a model of reactive astrocytes FA number and size were elevated along with the expression of plectin, highlighting involvement of plectin in pathological conditions.

## Introduction

Astrocytes play important roles in maintaining central nervous system (CNS) homeostasis by providing metabolic and neurotrophic support as well as modulating the plasticity and density of synapses in pathological conditions^1^. To this end, astrocytes rely on their morphological complexity, which include small processes that enable close contacts and bidirectional interactions with the microvasculature, other glial cells, and synapses. The structural integrity of these processes depends on cytoskeletal elements^2^.

In addition to microtubules (MTs) and the actomyosin system, astrocytes express a diverse intermediate filament (IF) network, composed of vimentin, glial fibrillary acidic protein (GFAP), synemin, nestin and nuclear lamins^2^. All these filamentous systems interact with each other and with other cellular structures, either directly, or indirectly through cytoskeletal linking proteins^3^. Plectin, a highly versatile cytolinker, is expressed widely in the CNS, with particularly high expression levels in astrocytes^4-6^. Alternatively spliced exons in the 5′-region of the *PLEC* gene result in a high diversity of plectin isoforms (P1, P1a – P1k)^7,8^, each with isoform-specific functions and subcellular targeting^9^. Mouse brain cortex astrocytes express predominantly isoforms P1c, P1e, and P1g, with P1c showing the highest expression level^6^. Though plectin research has mainly focused on the protein’s role in skin and muscle pathologies, findings of neuropathies in patients with plectinopathies suggest a significant role in CNS pathologies as well^5,8,10-12^. For example, P1c-deficient neurons exhibit tau accumulation on MTs, impaired neuritogenesis, and memory dysfunction^13^. In addition, we have recently shown that plectin deficiency impairs the collective migration of astrocytes and affects their volume regulation, indicating plectin’s potential involvement in glioblastoma and brain oedema^6^. These effects are likely mediated by focal adhesions (FAs), which regulate astrocyte morphology by linking the cytoskeleton with the extracellular matrix (ECM)^14^. Previously plectin isoform P1f was shown to mediate interactions of IFs with FAs^15-17^, though P1f is not expressed in astrocytes^6^. The continuous flux of assembling and disassembling of FAs, known as FA turnover, must be tightly regulated during the morphological remodelling of cells and cell migration^18^. FAs represent sites where signalling pathways regulating cell migration, adhesion, and survival are initiated^19^. These processes are crucial in physiological contexts, such as during neural development^20^, as well as in many pathologies, including glioblastoma formation and invasion^21^. In mature astrocytes, FAs mediate responses to various mechanical stimuli, such as disturbances in the volume of extracellular space in brain oedema or changes in the ECM stiffness associated with deposits of amyloid plaques in Alzheimer’s disease^22^. Following virtually any insults to the CNS, astrocytes undergo a process termed reactive astrogliosis, which involves morphological, molecular, and functional changes. Here, depending on the context, astrocytes may adopt multiple states, exhibiting a loss of some homeostatic functions and a gain of protective or detrimental functions simultaneously^23,24^. Reactive astrogliosis is frequently characterized by morphological remodelling and the upregulation of IFs, particularly vimentin and GFAP^24^. Given the wide spectrum of conditions that lead to reactive astrogliosis, and the distinct astrocyte phenotypes involved, there is a need for new molecular markers and functional readouts. In this study, we investigated the impact of plectin on modulation of FAs and its potential contribution to reactive astrogliosis and explored whether plectin could serve as a new molecular marker of this process.

We demonstrate that plectin is upregulated in conditions mimicking reactive astrogliosis and plays a critical role in regulating not only astrocyte morphology, FA dynamics and cytoskeletal organization but also significantly influences migration and the viscoelastic properties of single astrocytes. These results emphasize plectin’s importance in both, physiological processes like neurodevelopment, and pathological conditions leading to reactive astrogliosis.

## Methods

### Cell cultures

Experiments were performed on primary (*Plec^+/+^* and *Plec^-/-^*) and immortalized (*Plec^+/+^p53^−/−^* and *Plec^-/-^p53^−/−^*) mouse astrocytes. *Plec^+/+^* (wild-type) and *Plec^-/-^* (plectin-null) littermates were obtained by inter-crossing *Plec^+/−^* heterozygotes of the mouse strain Plec/29.B6 (N12)^25,26^ and *Plec^+/+^p53^−/−^* and *Plec^-/-^p53^−/−^* littermates by inter-crossing *Plec^+/−^p53^+/−^* heterozygotes of previously generated mouse line^26^. Both heterozygote mouse lines were maintained by backcrossing to a C57BL/6 background. Astrocytes were isolated from the cerebral cortices of neonatal mice following previously established protocols^27^ and cultured in standard cell culture medium consisting of high-glucose Dulbecco’s modified Eagle’s medium (DMEM, Thermo Fisher Scientific, Karlsruhe, Germany) supplemented with 10% fetal bovine serum, 1 mM sodium pyruvate, 2 mM L-glutamine, 5 U/ml penicillin, and 5 µg/ml streptomycin (DMEM+ medium). Astrocyte cultures were maintained in the incubator (37 °C, 5% CO_2_). Prior to experiments cell cultures were tested for astrocyte markers^6^.

Experiments were performed on cells plated on poly-D-lysine (1% v/v in dH_2_O) and laminin-coated (1% v/v in sterile PBS) coverslips. In a subset of experiments cells were washed twice with serum-free neurobasal medium (NB; Thermo Fisher Scientific, Waltham, MA, USA) and for 48 h maintained in NB+ medium consisting of NB complemented with B27 supplement (2%; Thermo Fisher Scientific), hbEGF (5 ng/ml), GlutaMAX (2 mM; Thermo Fisher Scientific), 5 U/ml penicillin, 5 μg/ml streptomycin).

If not indicated otherwise, cell media and reagents for all the experiments in this study were obtained from Sigma-Aldrich (Merck, Darmstadt, Germany).

### Immunolabeling

Astrocytes were labelled by immunocytochemistry protocol as described^28^. Briefly, to immunolabel vinculin, cells were washed with phosphate buffered saline (PBS), fixed in 4% formaldehyde (RT, 15 min) and permeabilized with 0.1 % Triton X-100 (RT, 10 min) (Merck Millipore, Germany). Immunolabeling of plectin, α-tubulin and vimentin was performed by dehydrating and permeabilizing astrocytes in ice-cold 100% methanol (10 min at -20 °C). Cells were then incubated in blocking solution (10% goat serum) for 1 h at 37 °C and then with mouse antibodies against vinculin (1:200; #ab18251, Sigma-Aldrich), rabbit antibodies against plectin (1:200; #46)^26^, antibodies against α-tubulin (1:1000; #ab18251, Abcam, Cambridge, UK) or goat anti-vimentin antisera (1:1000)^15^ for 2 h in the incubator (37 °C, 5% CO_2_) or overnight at 4 °C. Double immunolabeling was performed by applying antibodies sequentially. Secondary antibodies (goat anti-rabbit Alexa Fluor 488, goat anti-mouse Alexa Fluor 546, rabbit anti-goat Alexa Flour 546, 1:600; Invitrogen, Thermo Fisher Scientific, USA) were applied for 45 min at 37 °C. All antibodies were diluted in 3% BSA in PBS. Actin filaments were labeled in cells fixed with 4% formaldehyde (RT, 15 min) and incubated with 1× Phalloidin-iFluor 594 (Abcam, UK) diluted in 1% BSA (RT, 30 min). Coverslips were mounted onto glass slides using SlowFade Gold Antifade Mountant with 4′,6-diamidino-2-phenylindole (DAPI; Invitrogen, Carlsbad, CA, USA). For imaging samples with stimulated emission depletion (STED) microscopy secondary antibodies goat anti-mouse Abberior STAR ORANGE and goat anti-rabbit Abberior STAR RED were used (1:200, 45 min, 37 °C; Abberior Instruments, Germany) and mounted onto glass slides using anti-fade reagent (Mount Liquid Antifade, Abberior).

### Plasmids and transfection

Cells were transfected using plasmid DNA (pDNA) encoding for fusion proteins between and fluorescent proteins and various plectin isoforms (P1c2α3α-EGFP, P1e-EGFP, P1g-EGFP^29^), pDNA encoding for Vinculin-Venus (Addgene, plasmid #27300)^30^, and Paxillin-EGFP (Addgene, plasmid #15233)^31^. FuGENE 6 transfection reagent (Promega, USA) was used for pDNA transfection according to the manufacturer’s instructions. 48 h post-transfection live or immunolabeling experiments were conducted, as described above.

### Fluorescence microscopy

Imaging of fluorescently labeled samples was performed using the following microscopes: LSM 980 with Airyscan 2 (Carl Zeiss, Oberkochen, Germany), structured illumination microscope (SIM) ELYRA (Carl Zeiss) and STED microscope STEDYCON (Abberiror Instruments) combined with Zeiss Axio Observer 7 microscope (Carl Zeiss).

Unless stated otherwise, imaging of astrocytes with LSM 980 was conducted using a Plan-Apochromat 63×/1.4 oil objective (Carl Zeiss). Excitation was achieved using 405 nm, 488 nm, and 561 nm diode lasers. Emission fluorescence was filtered with 420-480 nm, 495-550 nm and 570-630 bandpass filters, respectively. During time-lapse imaging focus drift was compensated using Definite Focus module (Carl Zeiss). SIM microscopy was performed using an oil immersion objective (63×/NA 1.4, Carl Zeiss). Fluorophores were excited with 488 and 561 nm laser beams and emission fluorescence was filtered with 495–560 nm and 570–650 nm bandpass filters. Images were captured with an EMCCD camera (Andor iXon 885, Andor Technology, UK). To study the nanoscale organization of plectin in FAs we utilized STED microscopy. We used an oil-immersion plan apochromatic differential interference contrast (DIC) objective (100×, 1.46 NA, Carl Zeiss). Imaging was conducted using pulsed lasers at 561 nm and 640 nm for excitation and a depletion laser at 775 nm. The pixel dwell-time was set to 10 µs, and the pixel size was 20 nm × 20 nm. Laser power and detector or camera gain settings were optimized on control samples, devoid of any fluorophores.

### ELISA quantification

Primary mouse astrocytes were plated in triplicates on 24-well plates, with a seeding density of 50 × 10^4^ cells per well and incubated in DMEM+ or NB+ medium for an additional 48 h. Subsequently, the supernatant was collected and stored frozen at − 80 °C. Prior to performing indirect ELISAs, the protein concentration of samples was determined using a BCA protein assay (Thermo Fisher Scientific). Samples, diluted (1:1) in coating buffer (0.1 M Na2CO_3_/NaHCO_3_, pH 9.4), were loaded onto 96-well plates (Thermo Fisher Scientific), incubated for 2 h at RT, followed by washing (3×) in PBS/0.05% Tween-20, and incubated with blocking solution (washing solution plus 2% BSA) for 1 h at RT. The plates were then incubated (2 h, RT) with rabbit anti-plectin antiserum #9^26^(diluted 1:500 in blocking solution). For subsequent incubation with secondary antibodies, goat anti-rabbit HRP (Thermo Fisher Scientific) was used at a dilution of 1:4000. For signal detection, TMB substrate (Thermo Fisher Scientific) was added for 15 min and the reaction stopped with 2 M sulfuric acid. Immediately after, the absorbance was read at 450 nm using a Multiscan GO spectrophotometer (Thermo Fisher Scientific).

### Cell spreading assay

*Plec^+/+^*, *Plec^-/-^*, *Plec^+/+^p53^-/-^,* and *Plec^-/-^p53^-/-^* astrocytes were plated on coverslips as indicated above and allowed to spread for 30 min, 2 h or 5 h in the incubator (37 °C, 5% CO_2_). At each time point, cells were immunolabelled as described above. To observe the spread area and cell morphology, astrocytes were imaged using confocal microscope LSM980 (Carl Zeiss), equipped with DIC using a 20×/0.8 DICII objective.

### FA dynamics

To study FA turnover, astrocytes plated on cell culture tubes were transfected with pDNA encoding for Vinculin-Venus and Paxillin-EGFP. After 48 h cells were detached using trypsin-EDTA solution (Sigma, #T-3924) and plated on coverslips. Astrocytes were allowed to attach for 1 h in the incubator (37 °C, 5% CO_2_) and transferred to the preheated microscope stage. Standard cell culture medium was replaced by DMEM/F-12 medium without phenol-red (Gibco, #11039-021). Imaging was performed with LSM 980 with a 40×/1.4 oil objective (Carl Zeiss). FAs were recorded at the 3 min intervals for 2 h. To measure mobility of FA constituents (paxillin and vinculin, respectively), fluorescence recovery after photobleaching (FRAP) was utilized. Several FAs per one field-of-view were completely bleached using high intensity laser (set to 90% of the maximum laser power). Fluorescence recovery was recorded at 3 s intervals for a total duration of 5 minutes. For each FA, the time constant (τ) of fluorescence recovery and the percentage of fluorescence recovery were determined from fluorescence intensity profiles.

### Atomic Force Microscopy

Nanowizard II (JPK, Germany) was used in conjunction with an inverted optical microscope (Observer.Z1, Carl Zeiss). To determine the mechanical properties of astrocytes, probes with spherical tips were used (CONT-Silicon-SPM Sensor, spherical tip, 6.62 µm diameter, sQube, Germany). At the beginning of each experiment, the sensitivity of cantilever was determined on a clean glass coverslip in PBS, and the spring constant was calibrated using the thermal noise method^32^. Indentation was performed over an 8 × 8 grid (corresponding to 30 µm × 30 µm) above the cell nucleus and at the cell periphery, respectively. The force set-point was set to 1 nN, and the speed of indentation was kept at 2 µm/s to limit hydrodynamic effects. The sampling rate was set to 2048 Hz. Indentation (force-distance) curves were fitted using the Hertzian-Sneddon model for a spherical indenter in the JPK Data Processing software (JPK). Young’s modulus of cells was reported as the mean value of 64 measurements from each grid. During the recordings, cells were kept in the DMEM/F-12 medium inside the petri-dish heater at 37 °C for a maximum duration of 2 h.

To measure the viscous properties of isolated astrocytes, an extended delay in the constant-height mode of the nanoindentation procedure was set to 2 seconds. This extended delay allowed for the observation of cell relaxation dynamics. Changes in the vertical deflection of the cantilever resulting from cell relaxation were recorded and utilized to extract relaxation times, as described below.

### Data analysis

#### Analysis of distribution of FAs and colocalization of FAs with plectin and cytoskeleton elements

##### FA number and distribution

Following automatic thresholding using Otsu algorithm, FAs larger than 0.25 µm^2^ were counted using particle analyzer for Fiji^33^. The number of FAs was calculated for the peripheral and central cell regions, as follows. The peripheral region of the cell was obtained by first reducing the cell outline by 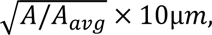 where *A* indicates the area of the particular cell and *A_avg_* represents the average cell area of the experimental group. Region between the cell outline and the reduced contour was designated as peripheral region, while the remaining portion of the cell was considered as central region.

##### Colocalization of FAs with plectin

The colocalization between immunolabelled FAs and plectin was measured using JACoP plugin^34^ in Fiji. Briefly, fluorescence micrographs were thresholded using Otsu algorithm (FAs larger than 0.25 µm^2^ were considered). Then, the degree of colocalization between FAs and plectin was calculated as the percentage of FA signal that is overlapped by plectin signal.

##### Colocalization measurements of FA with the cytoskeleton

The colocalization between FAs and cytoskeletal elements was measured using a custom-written Python script (Python Software Foundation, https://www.python.org/). Briefly, fluorescence micrographs (of immunolabelled FAs and respective types of cytoskeleton - microtubules, actin filaments, and vimentin) were thresholded using Otsu algorithm. Only FAs larger than 0.25 µm^2^ were considered. Colocalization of FAs and the cytoskeleton was then quantified as the percentage of vinculin signal overlapping with the signal from a specific cytoskeletal component. Moreover, the FA-cytoskeleton colocalization degree was calculated for the entire cells, peripheral and central regions separately.

##### Morphometry of astrocytes

Astrocytes were semi-automatically segmented based on the DIC and fluorescence (DAPI) images in Fiji. The cell area and the shape factor (4*π* ⋅ *A*/*P*^2^; where *A* represents the cell area and *P* the cell perimeter) were calculated for each cell.

#### Single cell migration

Live-cell imaging of single cell migration was performed 2 hours after cell plating using a Zeiss LSM980 confocal microscope equipped with a 20×/0.8 DICII objective and Definite Focus module. Multiple positions were selected on each coverslip to ensure representative sampling of the cell population. Time-lapse imaging was conducted over a period of 3 hours at an interval of 3 minutes. Post-acquisition, cells were manually tracked, and tracking data was processed using a custom Python script. Tracking coordinates (*x*, *y*) were converted to micrometres and centred at the origin for uniform analysis. From this data cell velocity, total displacement (cumulative path length) and maximal displacement (maximal distance between any two points on the track) were calculated.

#### Analysis of FA dynamics

##### FRAP

Fluorescence recovery curves were recorded by measuring the mean fluorescence intensity within the photobleached region over time. Original recordings were converted to relative units (with baseline representing 100%). These curves were fitted with a one-phase exponential model:

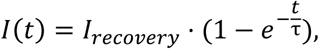

where *I* and *t* represent fluorescence intensity and time, respectively, whereas *τ* denotes the time-constant of the recovery. From the fitted curve, *τ* was obtained. Secondly, the percentage of mobile fraction was calculated as (*I_recovery_* − *I*_*post*_)/(*I_before_* − *I_post_*) × 100, here *I_before_*, *I_post_*, represents fluorescence intensity before and immediately after bleaching, whereas *I_recovery_* is the intensity after the return of the fluorescence to equilibrium. This model accounts for the rapid initial fluorescence recovery followed by a slower, asymptotic approach to equilibrium.

##### FA Turnover

FA turnover was analysed using TrackMate^35^, an open-source plugin for Fiji. Time-lapse images were loaded into TrackMate, and FAs were semi-automatically tracked over consecutive frames. The mean intensity of FAs was tracked over time from which assembly (formation) and disassembly (dissociation) rates of individual FAs were determined as follows: original data was log-linearized followed by linear regression to assembly and disassembly phases, respectively. The slope of fitted lines represents dissociation and association rates of FA. Only FAs with both phases present in experimental interval were included in the analysis.

#### Analysis of cytoskeleton connectivity and bundling

##### Connectivity

To analyse vimentin connectivity, we used a Python-based approach involving image processing and subsequent graph analysis. Images were pre-processed using Gaussian filtering and contrast enhancement (CLAHE) before skeletonization using the Otsu method. The skeletonized structures were converted into graph representations using the *skan* library^36^, allowing us to extract lengths of individual branches. We utilized NetworkX^37^ to build graphs of the filament network, identifying connected components, calculating degrees of nodes, and detecting endpoints. Data were visualized as plots of vimentin-network topology, providing insights into vimentin filament connectivity.

##### Bundling

The Gray Level Co-occurrence Matrix (GLCM) method was used to assess the texture of vimentin filament bundles by quantifying spatial relationships between pixel intensities. We utilized Python based-approach to compute GLCM features—correlation and dissimilarity, averaged over distances of 5 pixels and 4 angles (0, 45, 90, and 135 degrees). These features determine texture variations in filament bundling, allowing quantification of patterns like aggregation^38^.

#### Quantification of viscoelastic properties of astrocytes

Force-time curves, *F*(*t*), obtained using AFM, were split into indentation, relaxation, and retraction parts. Relaxation phase (see Fig. 7 for an example) was fitted using double exponential function for Maxwell viscoelastic body^39^:

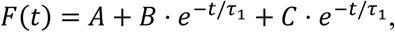

where *τ*_1_ and *τ*_2_ represent fast and slow relaxation times, respectively, while *A, B, C* are constants dependent on the elastic properties of the material. Maxwell body is a simple viscoelastic model that consists of a spring (representing elastic behaviour of a cell) and a dashpot (representing viscous behaviour of a cell) in series^39^.

#### Statistical analysis

All experiments were performed at least in a duplicate, on cells isolated from a minimum of two animals per genotype. The results are presented as the mean ± SEM. Statistical analysis was performed in SigmaPlot 11.0 (SYSTAT, USA). Statistical significance between two groups was determined with Student’s t test for normally distributed data or Mann-Whitney test otherwise. One-way ANOVA or ANOVA on ranks with Kruskal-Wallis post hoc test were used to test statistical differences between several groups. Statistical significances are considered as follows: \**P* < 0.05, \*\**P* < 0.01, \*\*\**P* < 0.001.

## Results

### In astrocytes plectin affects the integrity of FAs, leading to compromised single-cell migration

The morphological differentiation of astrocytes critically depends on FAs, which mediate interactions between the cells and the ECM. To investigate whether plectin colocalizes with FAs of astrocytes, we immunolabelled plectin and vinculin, a marker of FAs, and performed imaging with super-resolution microscopy (Fig. 1A). The results revealed that 48% ± 2% of the FA signal is overlapped by the plectin signal.

**Fig. 1.**
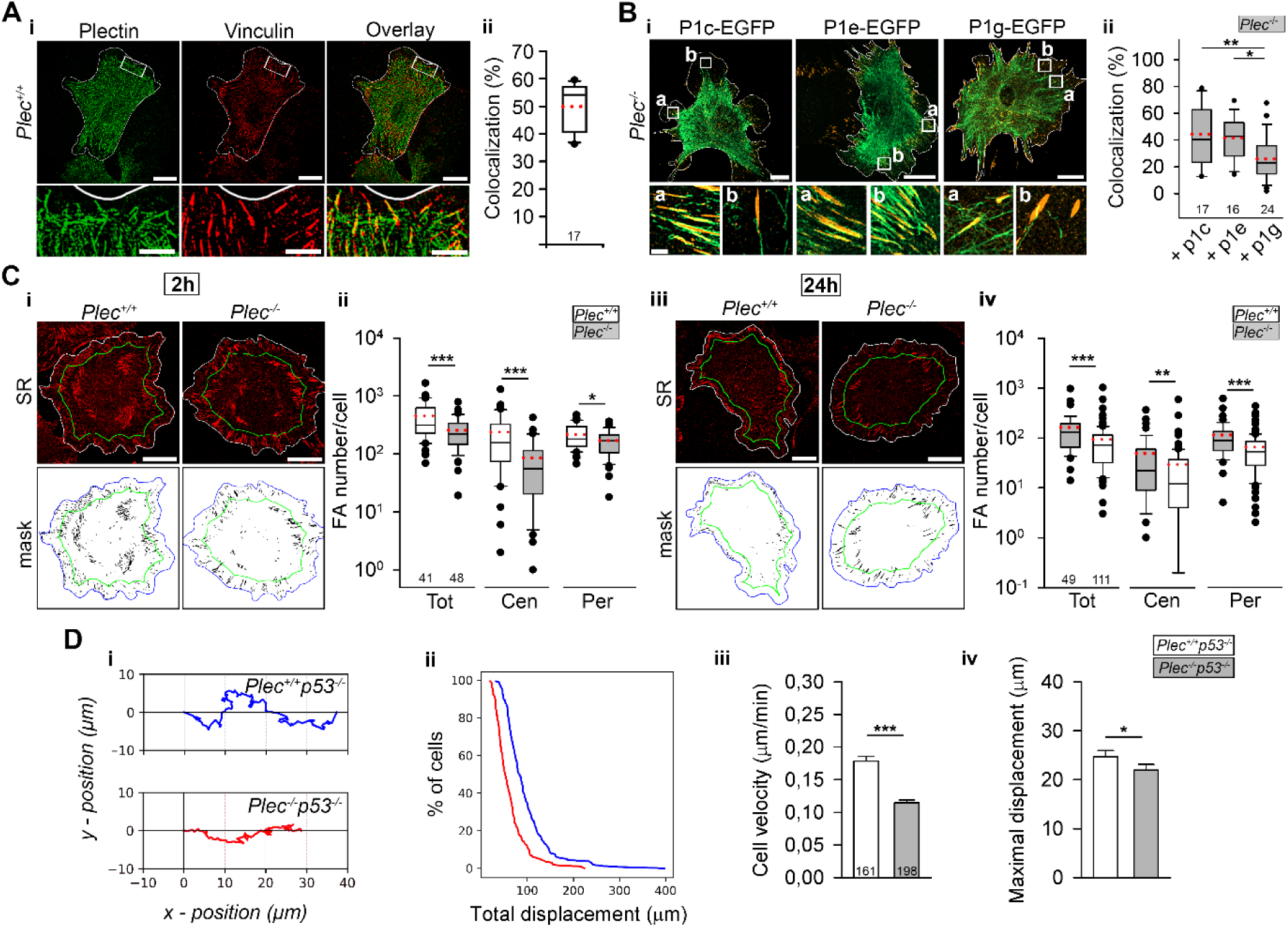
Plectin localizes with focal adhesions, determines their positioning and influences single-cell migration of mouse astrocytes. **Ai** Structured Illumination Microscopy micrographs of primary mouse astrocytes (*Plec^+/+^*), immunolabelled for plectin and vinculin. Below, enlarged regions of interest, the overlap of both signals shown on the right. Scalebars: 20 µm, 5 µm (inset). **Aii** ∼ 48% ± 2% of the FA signal is colocalized with the plectin signal. **B** Forced expression of EGFP-tagged plectin isoforms P1c, P1e, and P1g in plectin-deficient (*Plec^-/-^*) primary astrocytes immunolabeled for vinculin (red signal) (**Bi**). Below, enlarged regions of interest show the overlap of both signals (a, b). Scalebars: 30 µm, 2 µm (insets). **Bii** The percentages of overlapping signals for vinculin with P1c, P1e and P1g (P1c: 44% ± 5%, P1e: 42% ± 4%, P1g: 26% ± 3%; P1c vs. P1g, *P* = 0.005 and P1e vs. P1g, *P* = 0.02, One-Way ANOVA). **C** Super-resolution (SR) micrographs of *Plec^+/+^* and *Plec*^-/-^ astrocytes, immunolabelled for vinculin 2 h (**Ci**) and 24 h (**Ciii**) after plating. Cells are outlined in white (SR) or blue (masks), and green delineating central and peripheral regions. Scalebars = 30 µm. The number of FAs per region in *Plec^+/+^* and *Plec^-/-^* astrocytes after 2 hours (**Cii**): central region (Cen, 237 ± 39 vs. 85 ± 13, *P* < 0.001), periphery (Per, 216 ± 16 vs. 169 ± 12, *P* = 0.045), and entire cell (Tot, 453 ± 51 vs. 254 ± 23, *P* < 0.001), and after 24 hours (**Civ**): Tot (162 ± 23 vs. 94 ± 11, *P* < 0.001), Cen (49 ± 10 vs. 29 ± 6, *P* = 0.004), and Per (114 ± 14 vs. 65 ± 6, *P* < 0.001) (Mann-Whitney U test). **D** Representative tracks displaying migration of single immortalized (*p53^-/-^*) astrocytes expressing plectin (*Plec^+/+^*) or not expressing plectin (*Plec^-/-^*) over 3 hours (**Di**). The distribution of total cell displacement (**Dii**), mean cell velocity (**Diii**, 0.018 µm/s ± 0.008 µm/s vs. 0.011 µm/s ± 0.005 µm/s, *P* < 0.001, Mann-Whitney U test), and maximal cell displacement (**Div**, 25 µm ± 1 µm vs. 22 µm ± 1 µm, *P* = 0.02, Mann-Whitney U test) in *Plec^+/+^p53^-/-^* and *Plec^-/-^p53^-/-^* astrocytes. Black lines show the median and red dotted lines indicate the mean. The numbers below the boxplots indicate the number of cells analyzed.

To learn which plectin isoform mediates interactions with FAs, we transfected *Plec^-/-^* astrocytes with cDNAs expression plasmids of respective plectin isoforms (Fig. 1B). While the degree of colocalization between FA signal and either P1c or P1e signals was similar (44% ± 5% and 41% ± 4%, respectively), the colocalization of P1g with FAs was significantly lower (26% ± 3%). To study the effect of plectin deficiency on the number of FAs, *Plec^-/-^* astrocytes were cultured on laminin-coated coverslips for 2 h and 24 h (Fig. 1C). At 2 h post-plating, *Plec^-/-^* astrocytes exhibited significantly fewer FAs compared to *Plec^+/+^* astrocytes. This reduction was most pronounced in the central regions of astrocytes but less prominent at the cell periphery. 24 hours post-plating, the reduction in the number of FAs in *Plec^-/-^* astrocytes was noticeable across all cell regions. Similar effects were observed in *Plec^-/-^p53^-/-^* astrocytes, where forced-expression of P1c but not P1e or P1g partially rescued the reduced number of FAs and cell area (Fig. S1A-C).

Recently, we have shown that plectin critically affects the collective migration of astrocytes^6^. In the current study, single immortalized astrocytes were allowed to randomly migrate on laminin-coated coverslips for a total duration of 3 hours (Fig. 1D). Mean cell velocity, total cell displacement, and maximal displacement of *Plec^-/-^ p53^-/-^* were all significantly lower compared to *Plec^+/+^p53^-/-^* astrocytes. These results demonstrate the critical role of plectin in mechanisms controlling the mobility of single astrocytes, a property related to the physiological (neurodevelopment) as well as pathological conditions (glioblastoma).

FA formation begins with the assembly of nascent focal complexes at the cell membrane, followed by their maturation and disassembly. As FA mature, they recruit proteins, such as talin, paxillin, and vinculin, an adaptor protein essential for anchoring the actin cytoskeleton^18,40^. To elucidate the role of plectin in maturation of FAs, we utilized STED microscopy, where protrusions of astrocytes were imaged in cells immunolabelled for plectin and vinculin (Fig. 2A). This analysis revealed that in peripheral FAs, plectin signal is predominantly filament-like, while in centrally located FAs, plectin signals appear plaque-like that partially cover individual FAs (Fig. 2B and Fig. S2). To further assess the role of plectin in FA maturation, we quantified the degree of FA-plectin colocalization based on their distance from the leading edge. To this end, we first determined the principal axis of FA orientation, by averaging orientation of FAs in STED micrographs. Individual FAs were then projected onto this axis to analyse the correlation between distance from the cell edge and the extent of FA-plectin overlap (Fig. 2C). The localisation of plectin in FAs increases progressively from the cell edge within lamellipodia towards the cell centre, peaking at approximately 50 µm. Furthermore, plectin exhibits spatial polarisation within individual FAs, with preferential localisation at the tip of FA directed towards the cell edge (Fig. 2D). This spatial polarisation is most pronounced at 50 µm from the cell edge, aligning with the region of maximum colocalization (Fig. 2E). These results suggest that plectin plays an important role in the mechanism regulating FA maturation.

**Fig. 2.**
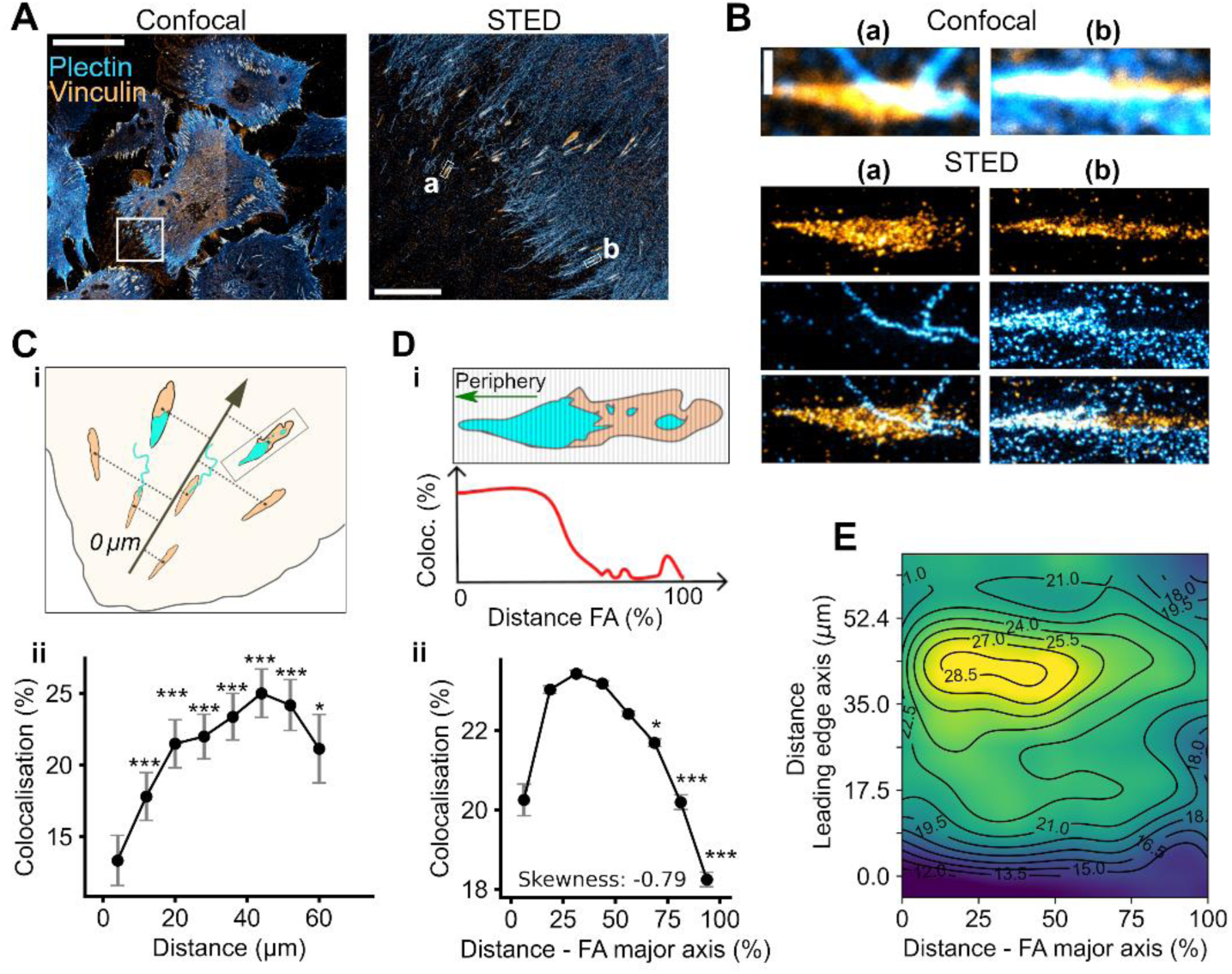
The distribution of plectin in focal adhesions is polarized and depends on the level of focal adhesion maturation. **A** Confocal micrograph of astrocytes immunolabelled for plectin (blue) and vinculin (orange). STED micrograph from the region framed in panel **A**. Insets show peripheral (a) and more central (b) focal adhesions (FAs). **B** Confocal and STED micrographs of vinculin and plectin signals of FAs from panel **A**. The plectin signal in the peripheral FA (a) is less abundant and filamentous, whereas the more central FA (b) shows a more abundant, diffuse plectin signal, primarily at the cell periphery-facing FA tip. **C** A schematic showing the mean orientation of all FAs, with an arrow pointing towards the cell centre (**Ci**). The distances of each FA from the most peripheral FA (assigned the distance of 0 µm), were determined by projecting them perpendicularly onto the mean orientation axis. The percentage of colocalization between plectin and vinculin increases with the distance from the leading edge (indicating FA maturation), peaking at ∼50 µm (**Cii**, \**P* < 0.05 and \*\*\**P* < 0.001, compared to the first point, Kruskal-Wallis test). **D** The degree of plectin localization in individual FA analyzed along the principal axis (**Di**), showing a preferential localization of plectin towards the FA tip, directed towards the cell periphery (**Dii**, \**P* < 0.05 and \*\*\**P* < 0.001, compared to the first point, Kruskal-Wallis test, Skewness = -0.79). **E** The colour-coded density plot of the dependency of plectin and vinculin colocalization (%) on the distance from the cell leading edge and the distance along the principal FA axis. Colocalization degree is color-coded, with contours indicating equal colocalization levels (yellow highlighting higher colocalization). Plectin polarization at the tip of FAs increases with distance from the leading edge and is peaking at ∼50 µm, coinciding with the highest degree of plectin colocalization in FA. *n* = 2075 FAs.

We next examined how plectin deficiency affects FA dynamics. FAs continuously interchange their constituents with the cytosol, undergoing turnover as part of mechanosensing and cellular mechanoresponses^41^. We used time-lapse confocal imaging to measure FA turnover, whereas Fluorescence Recovery After Photobleaching (FRAP) was utilized to measure the mobility of vinculin and paxillin in FAs. FA assembly and disassembly rates were determined from the time course of mean fluorescence intensity changes during the active spreading of astrocytes on laminin-coated substrates over a period of 2 h (Fig. 3A). Compared to wild-type (*Plec^+/+^*) astrocytes, plectin deficiency (*Plec^-/-^*) resulted in a significant decrease in FA assembly rate and an increase in the disassembly rate, resulting in a decreased assembly-to-disassembly ratio (Fig. 3B). Furthermore, we assessed the mobility of FA proteins vinculin and paxillin (Fig. 3C and 3D respectively). Complete photobleaching of individual FAs was followed by an exponential increase in the mean fluorescence intensity of the bleached region, indicating protein exchange between FA-bound (bleached) and cytosolic fractions. Compared to *Plec^+/+^* astrocytes, in *Plec^-/-^* astrocytes the characteristic recovery time (τ) was significantly longer whereas mobile fraction was not affected (Fig. 3C). In contrast, the absence of plectin did not regulate the speed of recovery for paxillin but increased its mobile fraction by 5% (Fig. 3D). These findings suggest that plectin is involved in regulating the maturation and turnover of FAs in astrocytes, as well as the kinetics of individual FA constituents, with the most pronounced effect observed on vinculin.

**Fig. 3.**
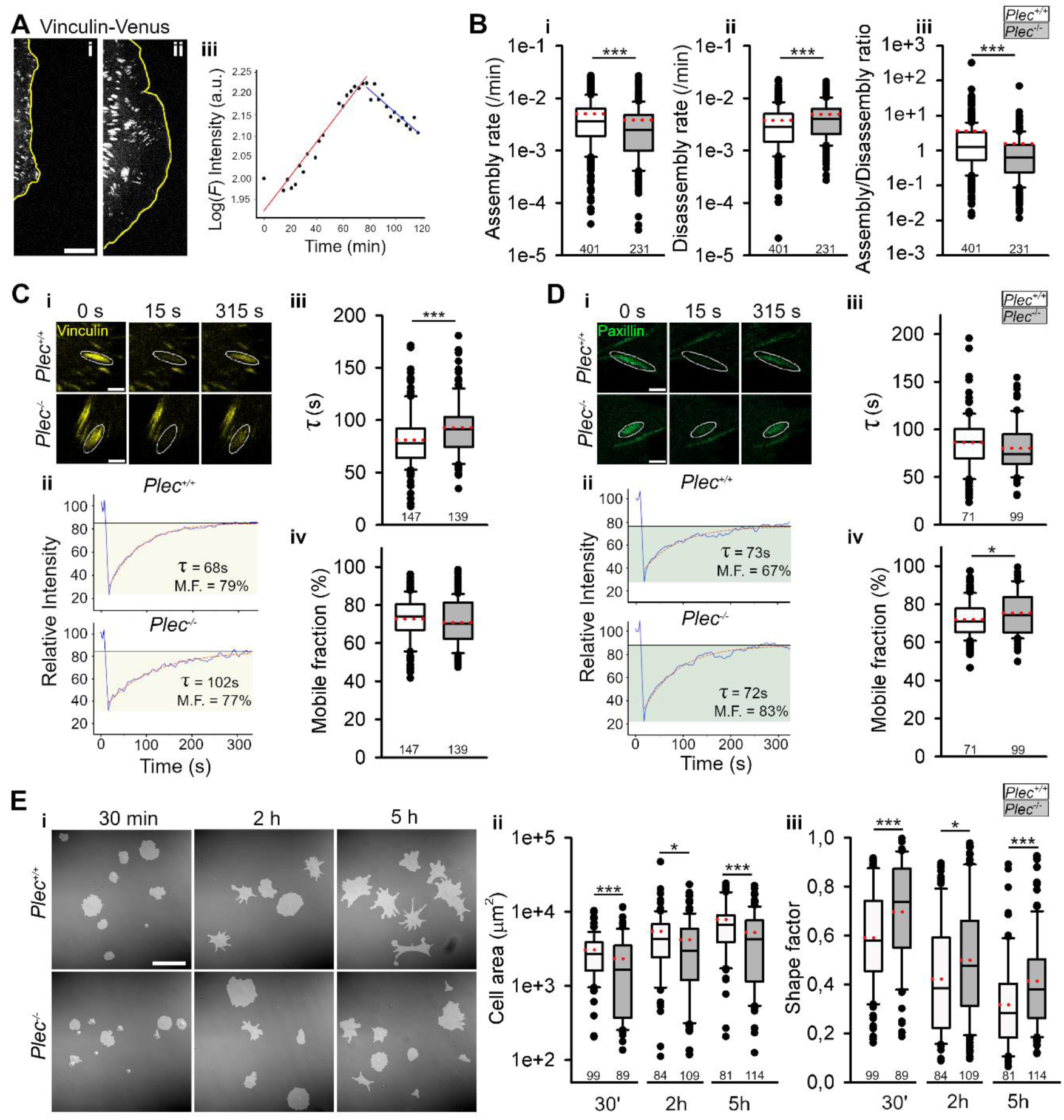
Plectin regulates focal adhesion turnover and constituent mobility leading to reduced cell spreading and a more circular morphology of astrocytes. **A** Focal adhesion (FA) turnover was monitored by forced expression of vinculin-Venus in primary plectin-expressing (*Plec^+/+^*) and plectin-deficient (*Plec^-/-^*) astrocytes. The leading edge of a spreading astrocyte at the start (**Ai**, 0 min) and end of imaging (**Aii**, 120 min). Scalebar = 20 µm. **Aiii** Time-dependent mean intensity of vinculin-Venus fluorescence in a single FA. Red and blue lines indicate least square fits for the assembly and disassembly phases, respectively. **B** The assembly (**Bi**, 5.0×10^-3^ ± 0.6×10^-3^ vs. 3.8×10^-3^ ± 0.5×10^-3^, *P* < 0.001) and disassembly (**Bii**, 3.8×10^-3^ ± 0.2×10^-3^ vs. 4.9×10^-3^ ± 0.3×10^-3^, *P* < 0.001) rates, and their ratio (**Biii**, 3.6 ± 0.8 vs. 1.6 ± 0.3, *P* < 0.001) in *Plec^+/+^* and *Plec^-/-^* astrocytes (Mann-Whitney U test). Mobility of vinculin (**C**) and paxillin (**D**) was determined by fluorescence recovery after photobleaching (FRAP) measurements in *Plec^+/+^* and *Plec^-/-^* astrocytes expressing vinculin-Venus and paxillin-EGFP, respectively (**Ci**, **Di**). Scalebar = 2 µm. Fluorescence intensity-versus-time plots of representative FAs (**Cii, Dii**). The blue line represents the mean intensity of a single FA, whereas the mobile fraction of vinculin and paxillin is shaded in yellow and green, respectively. Red dashed lines indicate the best-fit curve of the recovery phase. Scalebar = 2 µm. The time constant of recovery (τ) of vinculin (**Ciii**, 81 s ± 2 s vs. 93 s ± 2 s, *P* < 0.001, Mann-Whitney U test) and paxillin (**Diii**) in *Plec^+/+^* and *Plec^-/-^* astrocytes. The mobile fraction of vinculin (**Civ**) and paxillin (**Div**, 72 % ± 1 % vs. 75 % ± 1 %, *P* = 0.036, *t*-test) in *Plec^+/+^* and *Plec^-/-^* astrocytes. **E** DIC images of *Plec^+/+^* and *Plec^-/-^* astrocytes, overlayed with transparent masks (**Ei**). Cell area (**Eii**, 3000 µm^2^ ± 200 µm^2^ vs. 2300 µm^2^ ± 300 µm^2^, *P* < 0.001; 5500 µm^2^ ± 600 µm^2^ vs. 4200 µm^2^ ±400 µm^2^, *P* = 0.022; 7800 µm^2^ ± 600 µm^2^ vs. 5200 µm^2^ ± 400 µm^2^, *P* < 0.001) and shape factor (**Eiii**, 0.59 ± 0.02 vs. 0.70 ± 0.02, *P* < 0.001; 0.42 ± 0.02 vs. 0.50 ± 0.02, *P* = 0.026; 0.32 ± 0.02 vs. 0.41 ± 0.02, *P* < 0.001) of *Plec^+/+^* and *Plec^-/-^* astrocytes at 30 min, 2 h, and 5 h after plating, respectively (Mann-Whitney U test). Scalebar = 160 µm. Black lines show the median and red dotted lines indicate the mean. The numbers below the boxplots indicate the number of FAs analyzed.

We next assessed the role of plectin in morphological differentiation of astrocytes during spreading. Primary wild-type (*Plec^+/+^*) and plectin null (*Plec^-/-^*) astrocytes were plated on laminin-coated coverslips and allowed to spread for 30 min, 2 h, and 5 h, respectively (Fig. 3E). Compared to wild-type astrocytes, plectin deficient astrocytes (*Plec^-/-^*) had significantly lower surface area at all time-points and exhibited increased circularity. Similar effects of plectin deficiency were observed in immortalized mouse astrocytes cultured on both poly-D-lysine or laminin-coated substrates (*p53^-/-^*, Fig. S3).

### Plectin is involved in the fine-tuning of the intermediate filament network of astrocytes

To assess the importance of plectin in mediating interactions between actin filaments, microtubules, and vimentin filaments with individual FAs in astrocytes, we immunolabelled the respective cytoskeletal components and FAs in *Plec^+/+^p53^-/-^* and *Plec^-/-^p53^-/-^* astrocytes, as well as in primary *Plec^+/+^* and *Plec^-/-^* astrocytes. The interaction between the cytoskeleton and vinculin-labelled FAs was determined as the average percentage of FA area overlapped by all three main cytoskeleton types (Fig. 4) for the three regions of astrocytes – total, peripheral, and central. The anchoring of actin filaments (F-actin) at FAs provides mechanical support and links the ECM to the intracellular cytoskeleton^42^. Our study supported this notion, as we have observed a high degree of overlap between actin filaments and FAs (Fig. 4A). In centrally positioned FAs, the average percentage of FA overlapped by actin was 14 % lower in *Plec^-/-^p53^-/-^* astrocytes compared to *Plec^+/+^p53^-/-^* astrocytes. However, plectin did not affect colocalization of actin with peripheral FAs, nor did it affect colocalization between actin and FAs at the level of whole cells. We next focused on microtubules due to their role in regulating the turnover of adhesion sites^43^. In agreement, we observed that microtubules are detected in the vicinity of peripherally and centrally positioned FAs (Fig. 4B). Nonetheless, the percentage of overlap between microtubules and FAs is fourfold lower than that of actin filaments, and plectin deficiency did not alter the degree of colocalization. The overlap of vimentin filaments with FAs in immortalized astrocytes was comparable to that of microtubules (Fig. 4C). However, compared to *Plec^+/+^p53^-/-^* astrocytes, *Plec^-/-^p53^-/-^* astrocytes exhibited a significant reduction in the colocalization of vimentin with FAs at the whole cell level. This reduction was primarily due to the pronounced decrease in colocalization of vimentin with FAs in the central region. A similar effect was observed in primary astrocytes (Fig. 4D), which otherwise exhibited a fourfold increase in colocalization of vimentin with FAs compared to their immortalized counterparts. Yet the percentage of vimentin-FA overlap was significantly reduced in total, central and peripheral regions. In short, our results indicate that among the three major cytoskeletal systems, plectin in astrocytes primarily modulates the interaction between FAs and IFs. Hence, we next focused on examining the effect of plectin on global connectivity of vimentin filaments.

**Fig. 4.**
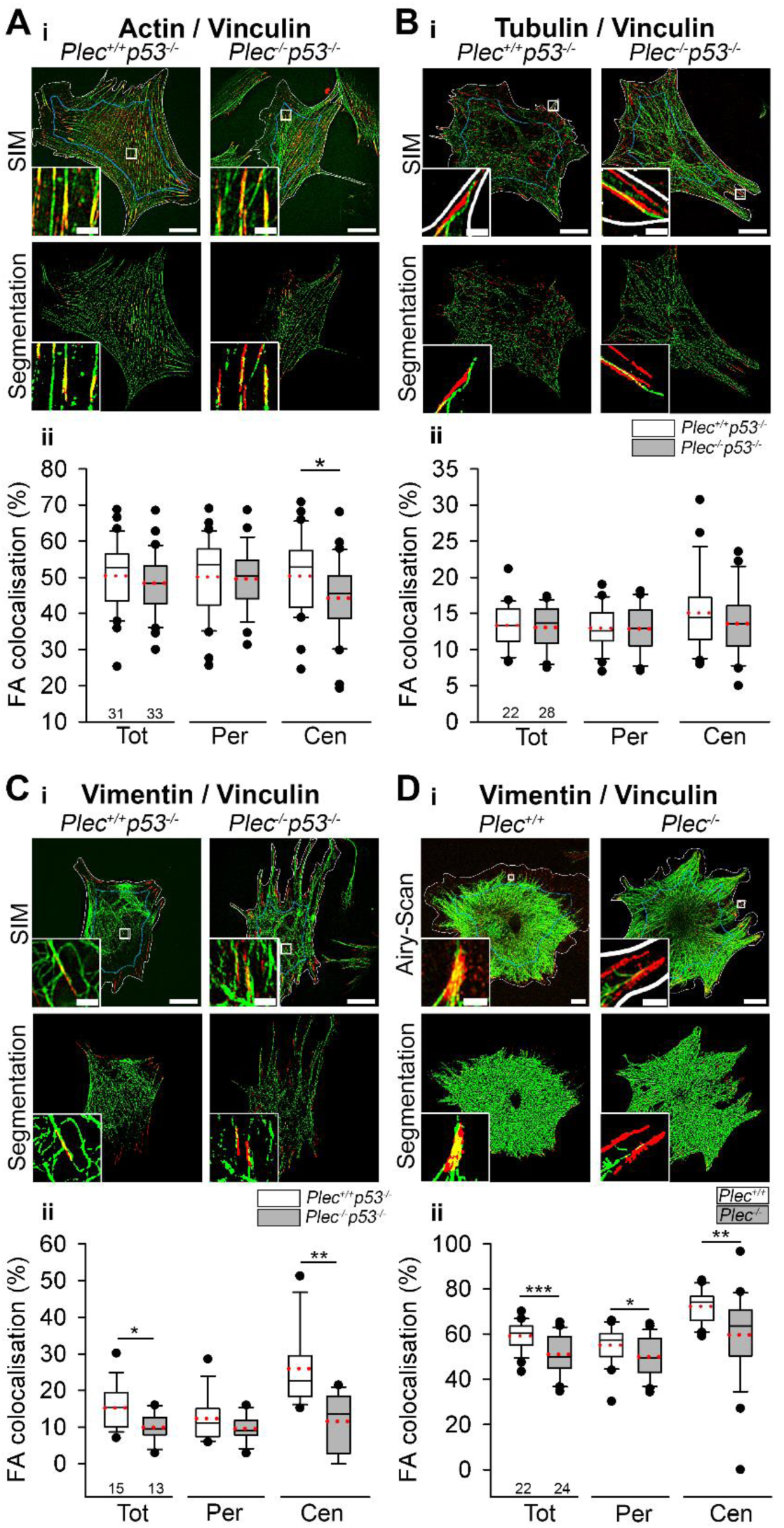
Plectin regulates the recruitment of actin and vimentin at focal adhesions in astrocytes. Colocalization of actin filaments (**A**), microtubules (**B**), and vimentin filaments (**C** and **D**) with focal adhesions (FAs) was quantified using super-resolution microscopy (structured illumination microscopy (SIM) or Airy-Scan). Immortalized (*p53^-/-^*, **A**–**C**) and primary astrocytes (**D**) expressing (*Plec^+/+^*), or not expressing plectin (*Plec^-/-^*) were investigated. Cell boundaries (white) define the total cell (Tot), with the blue line marking the separation between peripheral (Per) and central (Cen) regions, as shown in the upper rows. Bottom rows show segmented images. The overlap of cytoskeletal elements with FAs is visualized in yellow (**Ai-Di**, see enlarged insets). **A** The average percentage of FA signal overlapped by signal of actin filaments (**Aii**, Cen: 50% *±* 2% vs. 44*%* ± 2*%*, *P* = 0.028, *t*-test), microtubules (**Bii**) and vimentin (**Cii**, Tot: 15% ± 2% vs. 10% ± 1%, *P* = 0.010, *t*-test and Cen: 26% ± 3% vs. 12% ± 2%, *P* < 0.001, *t*-test) in *Plec^+/+^p53^-/-^* and *Plec^-/-^p53^-/-^* astrocytes in Cen, Per, and Tot regions. **Dii** The average percentage of FA signal overlapped by signal of vimentin in primary *Plec^+/+^* and *Plec^-/-^* astrocytes (Tot: 59% ± 1% vs. 51% ± 2%, *P* < 0.001, *t*-test, Per: 55% ± 2% vs. 50% ± 2%, *P* = 0.048, *t*-test, and Cen: 72% ± 2% vs. 60% ± 4%, *P* = 0.005, Mann-Whitney U Test). Scalebar: 20 µm; inset: 2 µm. Black lines show the median and red dotted lines indicate the mean. The numbers below the boxplots indicate the number of cells.

Vimentin forms a cytoplasmic network that enhances mechanical stability of cells^44^. In fibroblasts, plectin binds to this network, crosslinks it, and anchors it to other cytoskeletal components and the plasma membrane^45^. We immunolabelled vimentin in *Plec^+/+^p53^-/-^* and *Plec^-/-^p53^-/-^* astrocytes, followed by imaging using structured illumination microscopy (SIM). To provide insight into possible effects of plectin on organization of the vimentin network, SIM micrographs were subjected to connectivity analysis to quantify total vimentin length, average branch lengths, and the number of branching points (Fig. 5A and Methods section). We observed that in *Plec^-/-^p53^-/-^* astrocytes the empty spaces in the vimentin network are larger compared to their plectin-positive counterparts, leading to a more fragmented network (Fig. 5B). Specifically, in *Plec^-/-^p53^-/-^* astrocytes the total length of vimentin filaments was conserved, but the lengths of terminal branches (B1) and node-to-node (B2) branches were significantly longer in *Plec^-/-^p53^-/-^* compared to control *Plec^+/+^p53^-/-^* astrocytes (Fig. 5C). Moreover, *Plec^-/-^p53^-/-^* astrocytes exhibit significantly fewer vimentin fragments (B0) and B1 branches and the numbers of N1 and N3 branchpoints (i.e. terminal branchpoints and branchpoints connecting three branches, respectively) are decreased compared to *Plec^+/+^p53^-/-^* astrocytes. Thus, in the absence of plectin, the vimentin network undergoes structural remodelling, characterized by reduced numbers of branches and nodes (branchpoints) with concomitant elongation of individual vimentin branches. Additionally, plectin deficiency leads to an increased degree of vimentin filament bundling. The phenomenon can be seen as the spatial accumulation of pixels with similar fluorescence intensity values (Fig. 5D). To quantify the degree of vimentin bundling, we used analysis based on Grey-Level-Cooccurrence-Matrix (GLCM) (refer to Methods section for details). GLCM from *Plec^-/-^p53^-/-^* astrocytes exhibit more diagonal form than GLCM from *Plec^+/+^p53^-/-^* astrocytes, suggesting increased spatial bundling of the vimentin cytoskeleton. In *Plec^-/-^p53^-/-^* astrocytes the increased degree of bundling is indicated by higher GLCM correlation, and decreased GLCM dissimilarity indices compared to *Plec^+/+^p53^-/-^* astrocytes. Altogether, our results demonstrate that plectin regulates both the global vimentin connectivity and local properties of vimentin network, such as vimentin bundling, likely through direct filament interlinking.

**Fig. 5.**
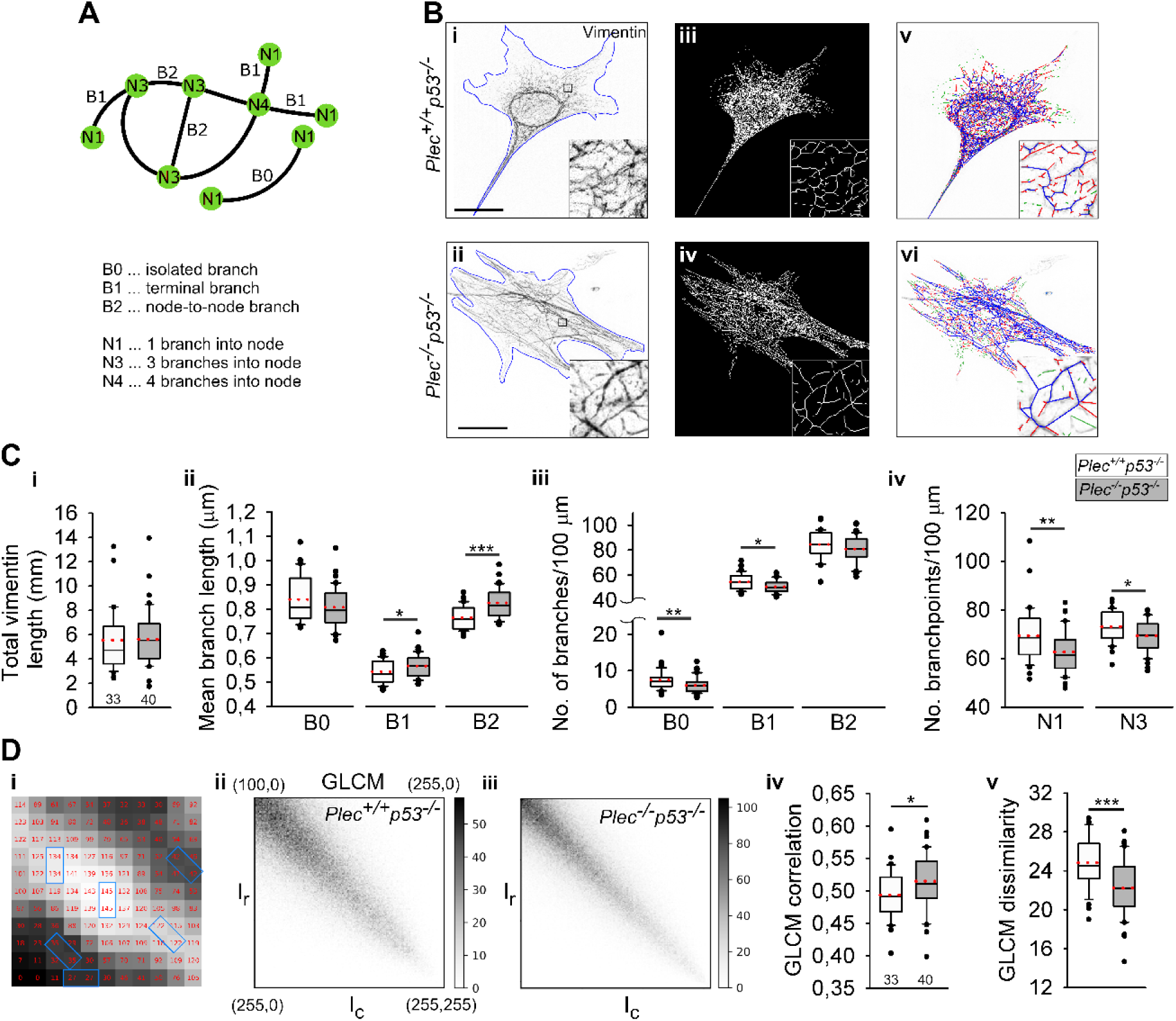
Plectin deficiency impairs vimentin network connectivity by increasing cytoskeletal branch length and decreasing the number of branches and branch points. **A** Connectivity analysis of branches (*B*) and branch points (*N*). **B** Structured illumination (SIM) micrographs of vimentin cytoskeleton in immortalized astrocytes expressing plectin (*Plec^+/+^p53^-/-^*) (**Bi**) or not expressing plectin (*Plec^-/-^p53^-/-^*) (**Bii**). Fluorescent signals were skeletonized (**Biii**, **Biv**) and subjected to connectivity analysis (**Bv, Bvi**) (see Methods). Branches of different types are colour-coded (B0 – green, B1-red, B2 – blue). **C** The total vimentin length (**Ci**) and mean branch length (**Cii**, B1: 542 nm ± 8 nm vs. 566 nm ± 7 nm, *P* = 0.03, B2: 765 nm ± 8 nm vs. 825 nm ± 9 nm, *P* < 0.001, *t*-test) in *Plec^+/+^p53^-/-^* and *Plec^-/-^ p53^-/-^* astrocytes. The number of branches (**Ciii**, B0: 7.4 ± 0.5 vs. 6.0 ± 0.3, *P* = 0.008, B1: 54 ± 1 vs. 50 ± 1, *P* = 0.011, both Mann-Whitney-U-test), and branchpoints (**Civ**, N1: 69 ± 2 vs. 63 ± 1, *P* = 0.007, Mann-Whitney-U test, and N3: 73 ± 1 vs. 69 ± 1, *P* = 0.017, *t*-test) per 100 µm in *Plec^+/+^p53^-/-^* and *Plec^-/-^p53^-/-^* astrocytes. **D** Texture analysis of the vimentin cytoskeleton in *Plec^+/+^p53^-/-^* and *Plec^-/-^p53^-/-^* astrocytes. **Di** displays an enlarged vimentin filament from a SIM micrograph, with red numbers depicting the fluorescence intensities of pixels. **Dii, iii** Grey-level-cooccurrence matrixes (GLCM) showing the comparison of neighbouring pixel intensities. In GLCM each element (*r - row*, *c - column*) represents the number of pixels with corresponding intensities (*I_r_, I_c_*); e.g., pixels with the same intensities, outlined in blue (**Di**), correspond to diagonal elements in the GLCM (**Dii**). GLCM from *Plec^-/-^p53^-/-^* astrocytes (**Diii**) show more diagonal form than GLCM from *Plec^+/+^p53^-/-^* astrocytes (**Dii**).GLCM correlation (**Dvi**, 0.052 ± 0.04 vs.0.050 ± 0.04, *P* = 0.034, *t*-test) and GLCM dissimilarity (**Dv**, 25 ± 3 vs. 22 ± 3, *P* < 0.001, *t*-test) in *Plec^+/+^p53^-/-^* and *Plec^-/-^p53^-/-^* astrocytes. Black lines show the median and red dotted lines indicate the mean. The numbers below the boxplots indicate the number of cells.

### Plectin regulates the viscoelastic properties of astrocytes

The organisation of the cytoskeleton, especially of IFs, is a critical determinant of cell stiffness^46^. Previous studies suggested that glial scars formed by reactive astrocytes can result in a softer mechanical environment, particularly due to ECM remodelling^47^. While mechanical properties of plectin deficient cells have been studied in non-neural cells^48^, these findings are not easily extrapolated to astrocytes due to differences in plectin isoform expression. Hence, we measured the mechanical properties of *Plec^+/+^* and *Plec^-/-^* astrocytes using atomic force microscopy (AFM). Individual cells were indented in the central part and at the cell periphery (Fig. 6A). The central region was divided into the nuclear (N), and the perinuclear (PN) parts. *Plec^-/-^* astrocytes exhibited decreased stiffness (lower Young’s modulus) at all three probed regions of the cell compared to *Plec^+/+^* astrocytes, most prominently above the cell nucleus and at the cell periphery (Fig. 6B). Cells are not purely elastic materials but exhibit viscoelastic (viscous and elastic) properties. To study viscous characteristics of astrocytes, the relaxation phase of the force-time nanoindentation data was fitted with two-exponent decay curve (Fig. 6C). Two relaxation times were extracted, τ_1_ and τ_2_, representing fast and slow relaxation components, respectively. τ_1_ originates from relaxation of the plasma membrane and cytosolic viscous gel-like properties, while τ_2_ is associated with the active reorganisation of the cytoskeleton^49^. *Plec^-/-^* astrocytes exhibited 11% slower τ_1_ relaxation time at the cell periphery in comparison to *Plec^+/+^* astrocytes. Conversely, plectin deficiency did not significantly affect the vicious properties above the cell nucleus or the slow relaxation component of viscosity. Thus, plectin deficiency impacts both the elastic and viscous properties of astrocytes, with the most pronounced effects observed at the cell periphery.

**Fig. 6.**
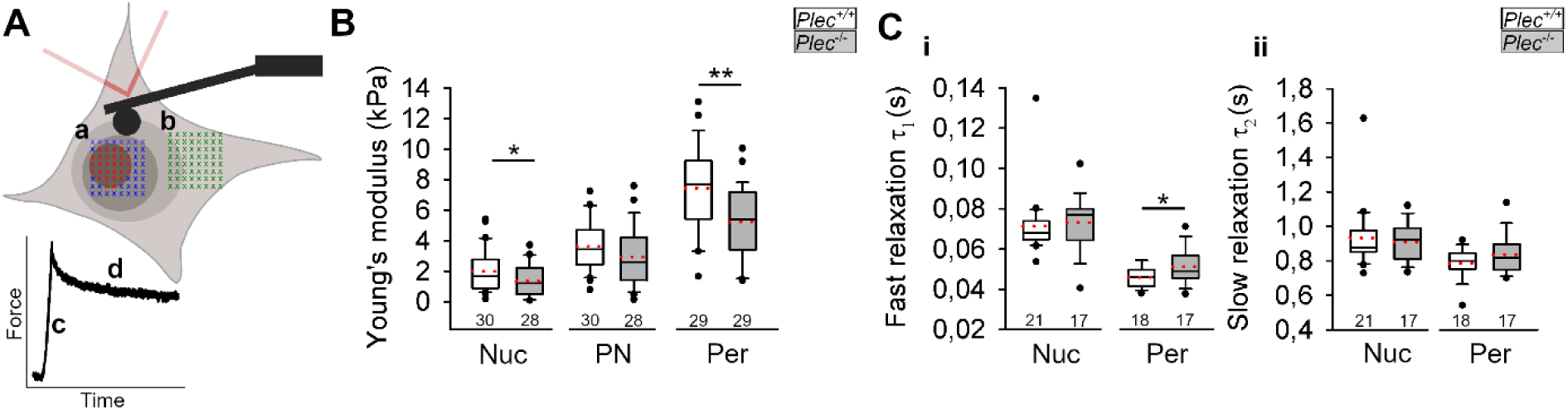
Plectin regulates the viscoelastic properties of mouse astrocytes. Young’s modulus and stress relaxation were measured using AFM. Astrocytes were mechanically probed in the central part of the cell (**Aa**) and above the periphery (Per, **Ab**). The central part of a cell was divided into nuclear (Nuc, red) and perinuclear (PN, blue) regions which were indented at multiple points. Below is a representative recording of a force-time curve with indentation (**Ac**) and relaxation (**Ad**) phase. The relaxation phase was fitted with a Maxwell model for viscoelastic materials to determine the fast (τ_1_) and the slow (τ_2_) components. **B** Young’s modulus of astrocytes, measured at Nuc, PN, and Per regions in primary astrocytes expressing plectin (*Plec^+/+^*) or not expressing plectin (*Plec^-/-^*) (Nuc: 2.0 kPa ± 0.3 kPa vs. 1.4 kPa ± 0.2 kPa, *P* = 0.044, Mann-Whitney U test, and Per: 7.4 kPa ± 0.5 kPa vs. 5.2 kPa ± 0.4 kPa, respectively, *P* = 0.002, *t*-test). **C** τ_1_ (**Ci**, Per: 0.046 s ± 0.001 s vs. 0.051 s ± 0.002 s, *P* = 0.04, *t*-test) and τ_2_ (**Cii**) in *Plec^+/+^* and *Plec^-/-^* astrocytes. Black lines show the median and red dotted lines indicate the mean. The numbers below the boxplots indicate the number of cells.

### Conditions mimicking reacitve astrogliosis lead to the upregulation of plectin and FAs

Hypertrophy of astrocytes, characterized by the enlargement of cell bodies and processes, and increased expression of IFs are hallmarks of reactive astrogliosis^24^. To gain insights into the role of plectin in morphological changes of astrocytes in reactive astrogliosis, we first assessed the morphometry of astrocytes by measuring the mean cell area. We compared morphometry parameters of astrocytes cultured in serum-free, neurobasal (NB+) medium (where astrocytes acquire native-like, i.e. stellate morphology)^50^, and of astrocytes cultured in serum-containing medium (DMEM+), which induces the phenotype of reactive astrocytes^50,51^. Our results support previous findings that DMEM+ astrocytes exhibit more pronounced hypertrophy, characterized by a threefold larger average cell surface area compared to NB+ astrocytes (Fig 7A). Reactive astrocytes are known to upregulate IFs^52^, actin and actin-related proteins^53,54^. In agreement, we observed upregulation of vinculin labelled FAs in DMEM+ vs. NB+ astrocytes. Quantitative analysis revealed both higher number and larger size of FAs in DMEM+ than NB+ astrocytes (Fig. 7B). Given that upregulation of IFs is a hallmark of reactive^55^, we hypothesized that also plectin would be upregulated. To this end, NB+ and DMEM+ astrocytes were immunolabelled with an antibody to plectin (Fig. 7C). Our results show that the total fluorescence signal of plectin in DMEM+ astrocytes was significantly higher compared to NB+ astrocytes. Consistently, mean intracellular plectin levels quantified using ELISA (Fig. 7D) were 48 % higher in DMEM+ astrocytes compared to NB+ astrocytes. These results suggest that plectin plays a pivotal role in morphological differentiation of astrocytes, which typically occurs during their maturation and in pathological conditions when astrocytes become reactive (Fig. 8).

**Fig. 7.**
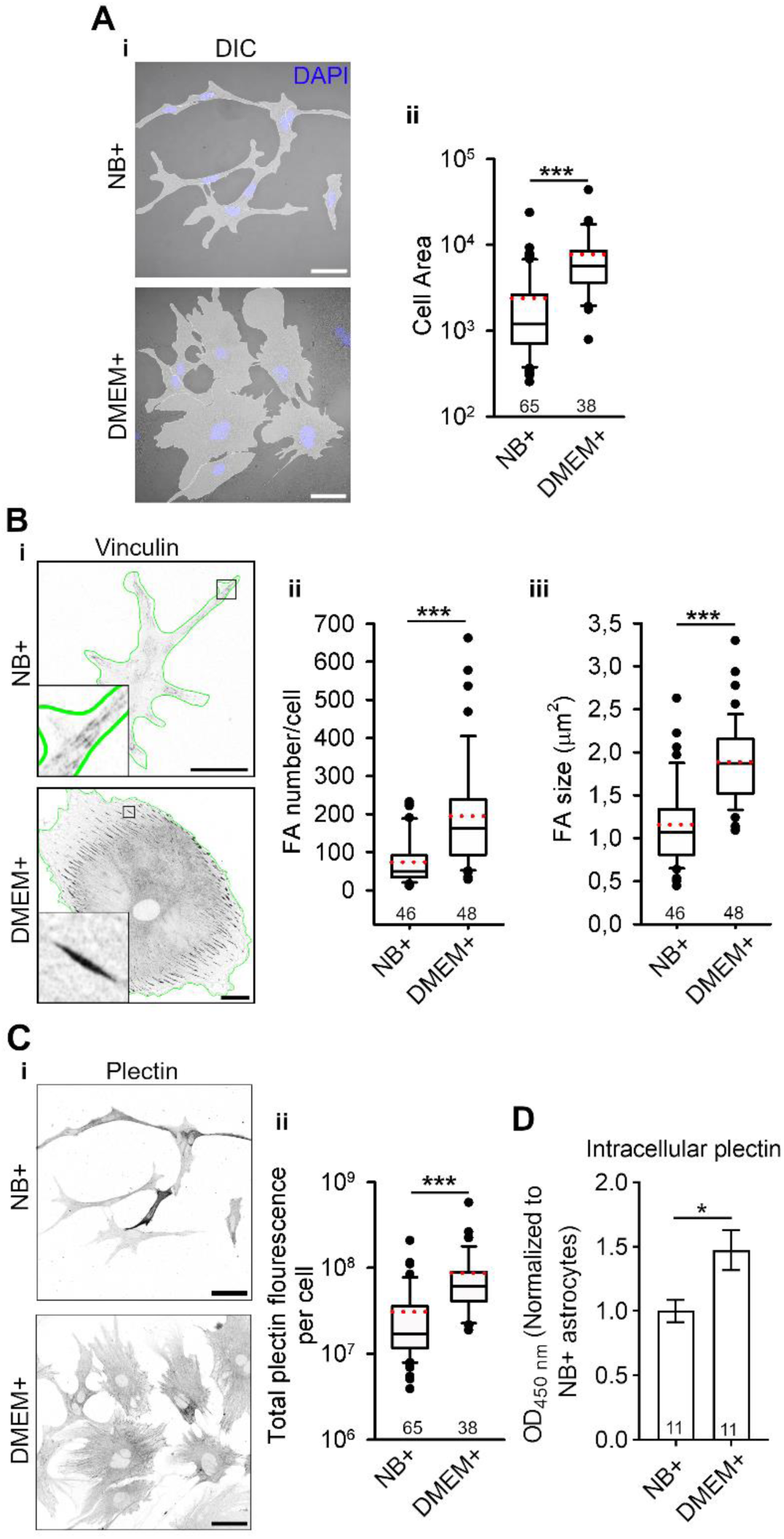
The number of focal adhesions and their size is increased in conditions mimicking reactive astrogliosis where plectin expression is elevated. **Ai** DIC micrographs overlayed with transparent masks of astrocytes grown in serum-free medium (NB+) and serum-containing culture medium (DMEM+). Scalebar = 50 µm. **Aii** Cell area of astrocytes grown in NB+ and DMEM+ medium (7749 µm^2^ ± 1215 µm^2^ vs. 2393 µm^2^ ± 426 µm^2^, *P* < 0.001, Mann-Whitney U test). **Bi** Confocal micrographs of vinculin-immunolabeled focal adhesions (FAs) in NB+ and DMEM+ astrocytes. Scalebar = 30 µm. Number of FAs per cell (**Bii**, 74 ± 9 vs. 195 ± 21, *P* < 0.001, Mann-Whitney U test) and FA size (**Biii**, 1.2 µm^2^ ± 0.1 µm^2^ vs. 1.8 µm^2^ ± 0.1 µm^2^, Mann-Whitney U test*, P* < 0.001) in NB+ and DMEM+ astrocytes. **Ci** Confocal micrographs depict the fluorescent signal of plectin in NB+ and DMEM+ astrocytes. Scalebar = 50 µm. **Cii** Total plectin fluorescence intensity per cell in NB+ and DMEM+ astrocytes (3.1×10^7^ a.u. ± 0.4 ×10^7^ a.u vs. 8.7×10^7^ a.u. ± 1.5×10^7^ a.u, *P* < 0.001, Mann-Whitney U test). **D** Intracellular plectin levels, measured by ELISA in NB+ and DMEM+ astrocytes (1.0 ± 0.1 vs. 1.5 ± 0.2, *t*-test, *P* = 0.015). Black lines show the median and red dotted lines indicate the mean. The numbers below the boxplots indicate the number of cells (A-C) or number of samples (D).

**Fig. 8.**
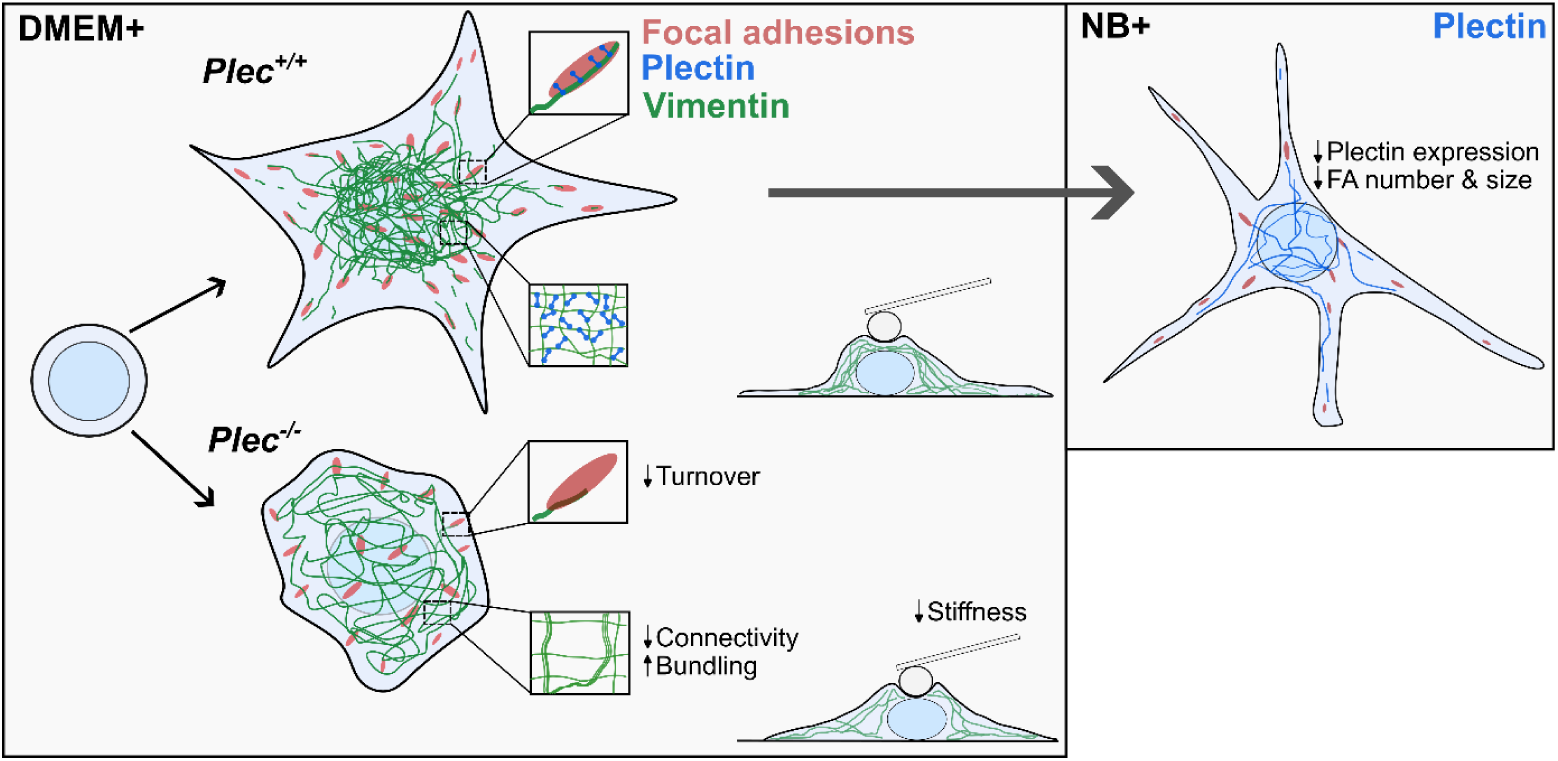
Schematic representation of the effects of plectin deficiency on focal adhesion-related processes in DMEM+ and NB+ astrocytes. DMEM+ cultured plectin-deficient (*Plec^−/−^*) primary astrocytes exhibit impaired cell spreading dynamics compared to wild-type (*Plec^+/+^*) cells, resulting in a more circular morphology and reduced cell surface area. This reduction in spreading is associated with fewer focal adhesions (FAs) in *Plec^−/−^* astrocytes and altered interactions of FAs with the cytoskeleton, particularly vimentin filaments. In *Plec^−/−^* astrocytes, the colocalization of vimentin with FAs is reduced (magnified boxed areas). Additionally, FAs in *Plec^−/−^* astrocytes demonstrate altered dynamics, specifically reduced turnover rates. The observed differences in cell morphology may also be driven by a general loss of vimentin network connectivity and filament bundling. These changes contribute to an overall softening of both the nuclear and peripheral regions of *Plec^−/−^* astrocytes. NB+ cultured mouse, which mimic in vivo-like physiological conditions, exhibit more pronounced stellation. NB+ astrocytes also show reduced expression of plectin, as well as fewer and smaller FA compared to DMEM+ astrocytes.

## Discussion

Our findings demonstrate that plectin is a key regulator of morphological differentiation of astrocytes as well as their mechanical properties, emphasizing its relevance to both physiological processes such as neurodevelopment and pathological conditions like reactive astrogliosis.

### Plectin Regulates FA Dynamics in Astrocytes

Our data demonstrate that in astrocytes plectin colocalizes with FAs and plays a significant role in regulating FA number and dynamics. Plectin-deficient astrocytes exhibit fewer FAs at 2 and 24 h after plating, a phenotype opposite to that seen in fibroblasts lacking plectin^15,56^. Unlike fibroblasts, which are mesodermal in origin, astrocytes are neuroectodermal-derived and have distinct developmental and functional characteristics that likely influences the composition of FAs and their associated interaction partners. Compared to fibroblasts, astrocytes express a distinct set of plectin isoforms. In particular, they do not express P1f which in fibroblasts specifically associates with FAs^15^. We show that all three isoforms colocalize with FAs to a similar degree in astrocytes. Notably, in immortalized *Plec^-/-^p53^-/-^* astrocytes, forced expression of isoform P1c isoform increases the number of FAs and partially restores decreased cell area, highlighting its role in FA stabilization and spreading of astrocytes.

While plectin has already been shown to play a role in collective astrocyte migration^6^, our study emphasizes its importance in single-cell migration. There are different requirements during collective versus single-cell migration, with plectin potentially facilitating stronger cell-cell and cell-ECM interactions in collective migration settings, while in single-cell migration it may play a key role in maintaining structural reorganization and cytoskeletal stability, ensuring effective and adaptive movement of individual cells^57^. In the latter case, *Plec^-/-^p53^-/-^* astrocytes displayed markedly reduced mobility. These findings align with the role of plectin in cytoskeletal organization and FA regulation which are both critical for maintaining directed and persistent migration^58^. Given the importance of astrocyte migration in neurodevelopment and the migration of glioblastoma cells (which can arise from neural stem cell-derived astrocytes^59^), the impaired migration observed in *Plec^-/-^p53^-/-^* astrocytes highlights the potential consequences of plectin dysfunction (or of its upregulation) in both physiological and pathological contexts.

During dynamic cellular processes such as spreading and migration, FAs undergo a complex maturation process that begins with the assembly of focal complexes near the leading edge. These nascent adhesions subsequently grow, elongate and recruit structural proteins like talin, paxillin, and vinculin, which stabilize the connection between the cell’s cytoskeleton and the ECM^60^. Our data reveal that near the leading edge of cells, plectin forms filament-like structures at FAs, suggesting its involvement in the early stages of FA formation. As FAs mature and shift toward the cell centre, plectin gradually covers part or all of the FAs. This aligns with findings from research on lung epithelial cells, where plectin predominantly localizes to central FAs and is typically absent from peripheral FAs^16^. The shift in plectin’s localization within FAs suggests that plectin may have distinct roles during FA maturation in astrocytes. These include supporting the dynamic assembly of nascent FAs at the periphery (filament like signal), potentially through interactions with cytoskeletal components like vimentin^15,61^, as well as stabilizing mature adhesions in central regions. Similar transitions in localization and content have been observed for other FA components, which exhibit spatial and functional shifts during FA maturation. E.g., newly formed focal complexes in lamellipodium are highly tyrosine phosphorylated, contain β_3_-integrin, talin, paxillin and low levels of vinculin and focal adhesion kinase (FAK), but are completely devoid of zyxin and tensin. The recruitment of these proteins into focal complexes occurs sequentially, so that their specific protein composition depends on their age^62^. Integrins also undergo a spatial shift, with β1 integrins becoming more prominent in central fibrillar adhesions during fibroblast spreading, as opposed to β3 integrins that dominate peripheral FAs^63^. Furthermore, we highlight a distinctive spatial polarization of plectin within individual FAs, where plectin predominantly localizes at the tips of FAs oriented towards the cell edge. Both observed distributions align with previous reports on the role of plectin in reinforcing cytoskeletal stability around the nucleus^15,17^. The progressive enrichment of plectin in FAs from the leading edge towards the cell centre underscores its role in regulating FA maturation, while its polarized localization in individual FAs suggests a specialized function in maintaining the directional dynamics of these structures. These findings provide new insights into the role of plectin in FA maturation and suggest its potential involvement in regulating cell adhesion and migration.

Our results demonstrate that plectin deficiency significantly affects FA turnover during astrocyte spreading by decreasing the assembly rate of FAs while conversely increasing their disassembly rate. This shift suggests that plectin is critical for maintaining a balance between the formation and breakdown of FAs. The decreased assembly rate in plectin-deficient cells may reflect a reduced ability to recruit or stabilize structural components necessary for the formation of nascent adhesions^64^. Whereas the increased disassembly rate implies a destabilization of existing FAs in the absence of plectin, further contributing to the overall decrease in FA stability. Conversely, plectin influences the kinetics of FA protein dynamics as observed through FRAP analysis of vinculin and paxillin. The longer recovery time in the absence of plectin suggests its compromised exchange between FAs and the cytosol. Vinculin interacts closely with the actin cytoskeleton and is regulated by cytoskeleton tension and adhesion turnover. In *Plec^-/-^* astrocytes, vimentin is known to reposition towards the cell periphery and form thick bundles near or in peripherally positioned FAs, which, in *Plec^+/+^* cells frequently interact with vimentin squiggles (short fragments of vimentin, roughly corresponding to B0 fragments in Fig. 5)^6,15^. This altered FA-vimentin interaction in plectin-deficient astrocytes may result in limited vinculin mobility. In contrast, paxillin’s recovery speed is not significantly affected by the absence of plectin. Its dynamics are less dependent on direct cytoskeletal linkages and more on signaling pathways^65^. As a result, the absence of plectin has little effect on paxillin turnover, leading to similar τ values in both *Plec^+/+^* and *Plec^-/-^* cells. However, the mobile fraction of paxillin slightly increases, indicating that plectin modulates the degree to which paxillin is exchanged between the FA and cytosolic pools. This indicates that plectin may influence the overall availability of paxillin in the dynamic exchange with FAs. This selective regulation of vinculin and paxillin kinetics underscores plectin’s multifaceted role in coordinating FA dynamics at the protein level. Collectively our results highlight that plectin plays a crucial role in maintaining the balance between the assembly and disassembly of FAs, as well as in regulating the kinetics of FA-associated proteins.

The importance of plectin in the morphological differentiation of astrocytes is further supported by the spreading assay, revealing that plectin-deficient astrocytes exhibit decreased cell surface area and increased circularity (less arborized morphology) compared to wild-type cells. It has been shown for several cell types that plectin serves as a multifunctional cytoskeletal crosslinker, including the anchoring of the cytoskeleton to membrane-associated structures such as FAs and adherens junctions ^8,45^. Our findings demonstrate that plectin regulates the recruitment of actin and vimentin to FAs in astrocytes, suggesting that plectin promotes the morphological differentiation of astrocytes by coordinating the interactions among FAs, actin, and IFs.

### Plectin primarily modulates vimentin cytoskeleton and is vital for astrocytic viscoelastic properties

Our study demonstrates the critical role of plectin in modulating the interactions between FAs and various components of the cytoskeleton in astrocytes. The strong overlap observed between actin filaments and FAs aligns with the fact that actin filaments dock in FAs. However, the decrease in actin-FA overlap in centrally positioned FAs of plectin-deficient astrocytes suggests that plectin facilitates the stabilization of actin-FA linkage specifically in this region. This finding aligns with the differential localization of plectin within FAs depending on their cellular position and supports our conclusion that plectin may play distinct roles during the maturation of these structures. Moreover, the vimentin-FA interaction is markedly affected by plectin deficiency, with significant reductions in colocalization in both immortalized and primary astrocytes, particularly in their central regions. This phenomenon has also been described in other cell types, such as fibroblasts, where FA-associated P1f anchors cage-like vimentin network surrounding the nucleus to the extracellular matrix^15^. Vimentin forms an intricate network throughout the cytoplasm that enhances mechanical stability, distributes stress, and provides resilience to cells^44^. Our connectivity analysis of the vimentin network demonstrates that plectin deficiency leads to a less dense vimentin network, elongated terminal branches and reduced branching points. These findings suggest that plectin regulates the dynamic interactions between cytoskeletal components and FAs and maintains the overall organization and mechanical stability of the vimentin network in astrocytes. This integrated network is crucial for structural support and resilience against mechanical stress, indicating that plectin is essential for the functionality and integrity of astrocytes under various (patho)physiological conditions. Of note, in astrocytes, vimentin is the most common co-polymerization partner of GFAP^66^ and expression of both vimentin and GFAP increase to a similar extent in reactive astrocytes^67^. As the interaction between plectin and GFAP has been validated using electron microscopy, immunofluorescence, and coimmunoprecipitation^4,11,68^, it is reasonable to infer that the results of our vimentin connectivity analysis would largely parallel those for GFAP.

Increased GFAP expression was shown to be accompanied by a softening of non-nuclear regions in reactive astrocytes^47^. Our findings in *Plec^-/-^* astrocytes show similar phenotypic shifts, including the loss of arborized morphology and softening of peripheral regions of astrocytes. Peripheral regions may *in vivo* correspond to astrocytic endfeet. These findings suggest that plectin plays a role in tuning the mechanical properties of astrocytes, specifically in regions that are vital for regulating the blood-brain barrier, cerebral blood flow, nutrient uptake, waste clearance, and integrating and modulating synaptic transmission. The mechanical properties of astrocytic peripheral regions are critical as they may regulate cerebral blood flow^69^, formation of tripartite synapses essential for cognitive functions^70^, and regulation of glymphatic flow^71^. Our results are in line with the softening of non-nuclear regions observed in reactive astrocytes due to the upregulation of IFs^47,72^. In plectin-deficient astrocytes, the vimentin cage is no longer confined to the perinuclear region^6^, with vimentin filaments extending more prominently toward the cell periphery. Hence, extension of vimentin network to the cell periphery in the absence of plectin likely contributes to the observed softening of peripheral cell regions. In agreement, astrocytes expressing plectin had more protrusions and were less circular, compared to the more rounded plectin deficient cells.

### Plectin is a key factor in morphological differentiation of astrocytes

The morphological plasticity of astrocytes is crucial for their function, especially during reactive gliosis, a process triggered in response to CNS injury or disease^24^. Reactive astrogliosis is characterized by astrocyte hypertrophy along with changes in cytoskeletal protein expression and alterations in cell shape^23^, and the formation of cell-matrix contacts like FAs^55^. Since these events are plectin-dependent, targeting plectin may provide new ways of modulating astrocyte responses during CNS pathologies. Our results are in line with previous reports demonstrating that astrocytes cultured in serum-containing medium (DMEM+) exhibit significant hypertrophy and a reactive phenotype in contrast to serum-free (NB+) medium-cultured astrocytes, which adopt a more native-like stellate morphology^50^. Here we report that FAs number and their size are higher in DMEM+ compared NB+ astrocytes, which aligns with increased F-actin and alpha actinin in reactive astrocytes^53^. Additionally, we show here that plectin is significantly upregulated in reactive (DMEM+) astrocytes compared to native-like state-resembling NB+ astrocytes. This suggests that plectin plays a pivotal role in cytoskeletal reorganization during reactive astrogliosis and is aligned with the upregulation of IFs, plectin’s major binding partners^2^. This upregulation of plectin likely contributes to the cytoskeletal changes necessary for the morphological and functional adaptations of astrocytes during reactive gliosis.

Overall, our findings demonstrate the multifaceted role of plectin in regulating cytoskeleton-dependent processes in astrocytes - from morphological differentiation and FA dynamics to cytoskeletal organization, migration, and cell mechanics. By coordinating the interplay between IFs, actin filaments and FAs, plectin enables astrocytes to adapt their structure and function during physiological processes and in response to pathological stimuli. Thus, our study advances not only our understanding the role of plectin in astrocyte biology, but also identifies plectin as a potential therapeutic target for conditions characterized by abnormal astrocyte behavior, such as during neurodegenerative diseases and in glioblastoma.

## Statements and declarations

### Funding

This work was supported by the Slovenian Research Agency (grant numbers P3-0310, P3-0083, J7-3153), the A4L_ACTIONS and A4L_BRIDGE projects funded by the European Union’s Horizon 2020 (grant agreements no. 964997 and 101136453), EuroCellNet COST Action (grant number CA15214), Network of infrastructure Centres of University of Ljubljana (I0-0022, I0-0048 CIPKeBiP, I0-0034 Celica), EU Interreg Italia-Slovenia Immunocluster-2.

### Competing interests

The authors declare no conflict of interest.

### Author contributions

B.F. performed experiments, analyzed data, prepared figures, and wrote the manuscript. M.P. assisted with the study conception and design and co-wrote the manuscript. V.M.P.D. performed experiments, analyzed data, prepared figures, and co-wrote the manuscript. R.Z., G.W. co-wrote the manuscript. J.J. designed and directed the study, prepared figures and co-wrote the manuscript. All the authors commented on previous versions of the manuscript and approved the final manuscript.

## Acknowledgements

The authors gratefully acknowledge Primož Runovc for his help with the experiments, prof. dr. Simon Horvat and dr. Urška Hostnik for the development and implementation of a rapid genotyping test for neonatal F2 crossbred offspring, and Katja Skulj for the maintenance and breeding of transgenic Plectin knockout lines.

## Data availability

All data that support the findings of this study are published in the article and are available from the corresponding author upon reasonable request.

## Ethics approval

The care of experimental animals and euthanasia of animals were performed in accordance with the following ethical codes and directives: International Guiding Principles for Biomedical Research Involving Animals developed by the Council for International Organizations of Medical Sciences and the Directive on Conditions for Issue of License for Animal Experiments for Scientific Research Purposes (Official Gazette of the RS, No.38/2013 and official consolidated text 21/18, 92/20, 159/21). The protocol for the euthanasia of animals used in our study was approved by the Veterinary Administration of the Ministry for Agriculture and the Environment of the Republic of Slovenia (permit numbers U34401-30/2021/8, U34401-27/2020/6, U34401-26/2020/4).

## Supplementary figures and figure legends

**Fig. S1.**
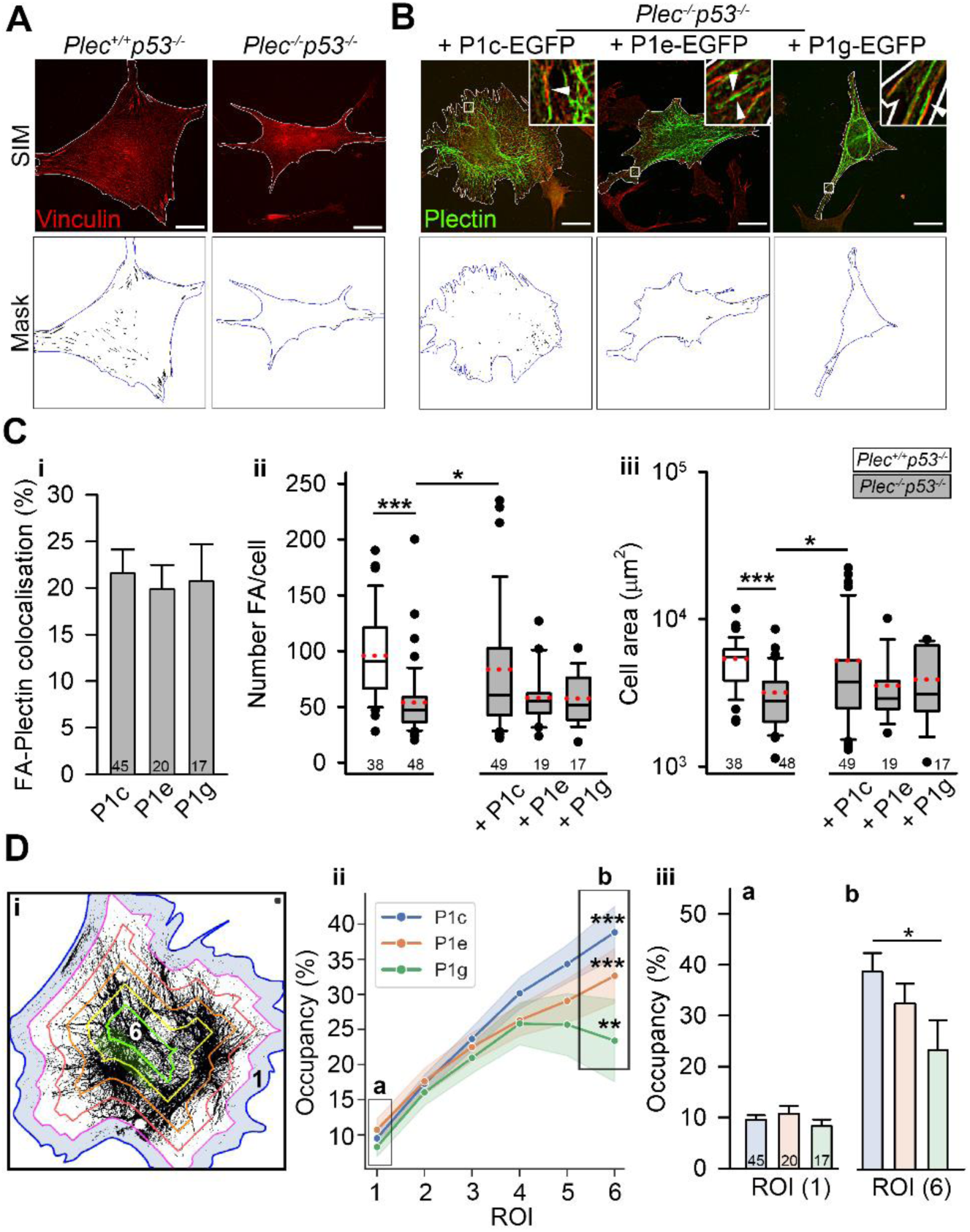
The forced expression of plectin isoforms in *Plec^-/-^p53^-/-^* astrocytes reveals focal adhesion upregulation by P1c plectin isoform. **A** Structured illumination micrographs of immortalized astrocytes expressing plectin (*Plec^+/+^p53^-/-^*) or not expressing plectin (*Plec^-/-^p53^-/-^*), immunolabeled for vinculin. **B** Forced expression of EGFP-tagged P1c, P1e, P1g plectin isoforms (green) in *Plec^-/-^p53^-/-^* astrocytes reveals plectin signal near or overlapping with vinculin-immunolabeled focal adhesions (FAs, red), as shown in the enlarged insets. **Ci** The percentage of colocalization between FAs and respective plectin isoforms (P1c, P1e, and P1g) **Cii** The number of FAs in *Plec^+/+^p53^-/-^* and *Plec^-/-^p53^-/-^* astrocytes (96 ± 6 vs. 54 ± 4, *P* < 0.001, Kruskal-Wallis test) and following forced expression of respective plectin isoforms in *Plec^-/-^p53^-/-^* astrocytes (P1c vs. *Plec^-/-^p53^-/-^*: 83 ± 10 vs. 54 ± 4, *P* = 0.02, Kruskal-Wallis test). **Ciii** The cell area of *Plec^+/+^p53^-/-^* and *Plec^-/-^p53^-/-^* astrocytes (5400 µm^2^ ± 300 µm^2^ vs. 3200 µm^2^ ± 200 µm^2^, *P* < 0.001, Kruskal-Wallis test), and of *Plec^-/-^p53^-/-^* astrocytes following forced expression of respective plectin isoforms (P1c vs. *Plec^-/-^p53^-/-^*: 5200 µm^2^ ± 700 µm^2^ vs. 3200 µm^2^ ± 200 µm^2^, *P* = 0.04, Kruskal-Wallis test). **Di** Plectin distribution in astrocytes, divided into 6 equally spaced regions of interest (ROIs). Subplasmalemmal (1) and central (6) regions of interest (ROIs) are highlighted in blue and green, respectively. **Dii** Occupancy of respective plectin isoforms in consecutive ROIs. Occupancy in ROI 1 (a) vs. ROI 6 (b) (P1c: 10% ± 1% vs. 39% ± 4%, *P* < 0.001, Paired t-test; P1e: 11% ± 2% vs. 33% ± 4%, *P* < 0.001, Paired t-test; P1g: 8% ± 1% vs. 23% ± 6%, *P* = 0.007, Wilcoxon Signed Rank Test). **Diii** Comparison of occupancy between the three isoforms in ROI 1 (a) and ROI 6 (b) (P1c vs. P1g: 39% ± 4% vs. 23% ± 6%, *P* = 0.039, Kruskal-Wallis test). Scalebar = 20 µm. Medians are shown as solid black lines, means as red dots. Numbers below boxplots indicate the number of cells.

**Fig. S2.**
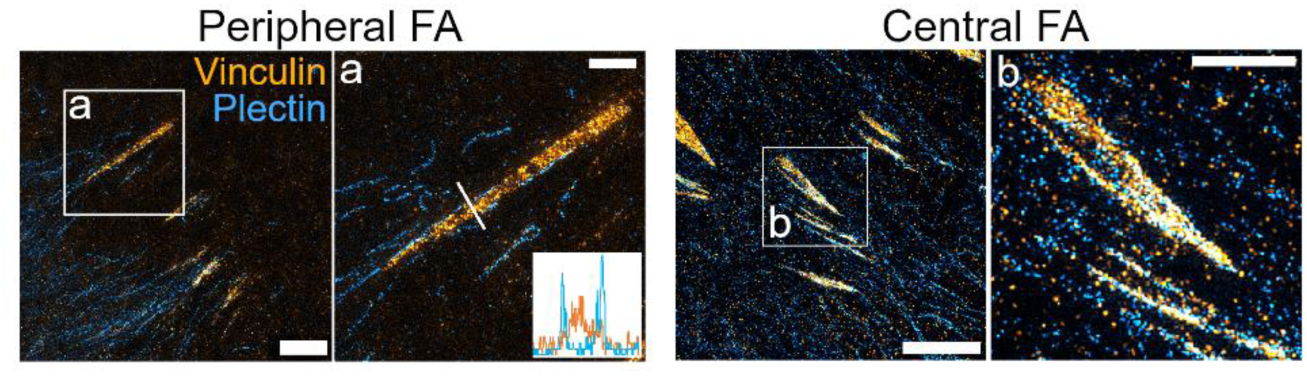
Focal adhesions and plectin exhibit distinct interaction patterns at the cell periphery compared to the central regions of the cell. Stimulated emission depletion microscopy (STED) micrographs of astrocytes immunolabelled for plectin (blue) and vinculin (orange). In peripheral focal adhesions (FAs) (**a**), plectin was predominantly positioned adjacent to FAs (note line profile in the enlarged boxed area), whereas in central FAs plectin typically covered a portion or the whole area of individual FAs (**b**). Scalebars: 5 µm and 2 µm (magnified areas).

**Fig. S3.**
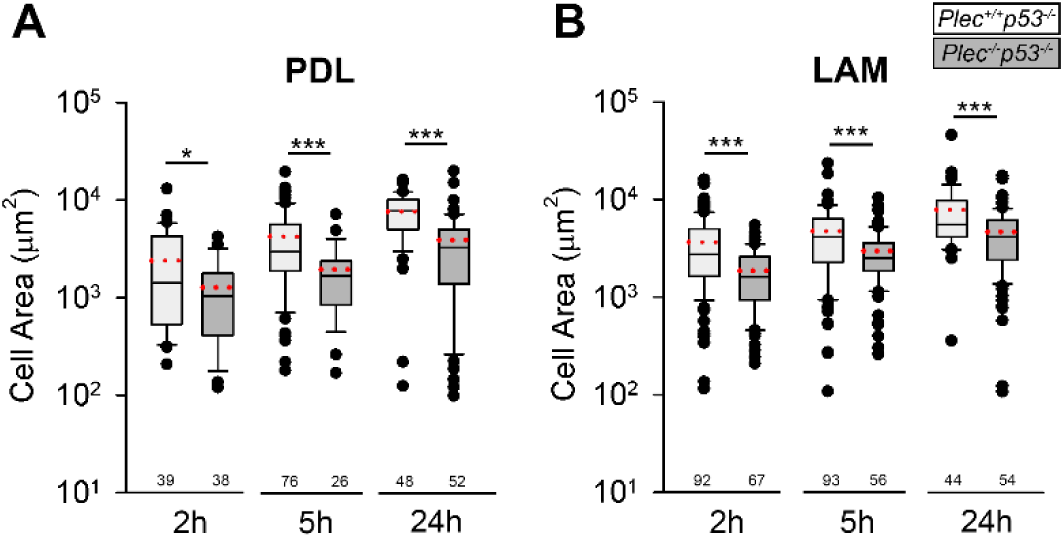
Plectin deficiency attenuates cell spreading of immortalized mouse astrocytes on both poly-D-lysine (PDL)- and laminin-coated substrates. Immortalized mouse astrocytes expressing plectin (*Plec^+/+^p53^-/^*^-^) or not expressing plectin (*Plec^-/-^p53^-/^*^-^) were plated on glass coverslips coated with either PDL (**A**) or laminin (**B**) and allowed to spread for 2 h, 5 h, or 24 h. Graphs show the cell area of *Plec^+/+^p53^-/^*^-^ and *Plec^-/-^p53^-/-^* astrocytes across all time points and on both substrate coatings. Numbers below boxplots indicate the number of cells. \**P* < 0.05, \*\*\**P* < 0.001 (Mann-Whitney U test). PDL – Poly-D-Lysine, LAM – Laminin.

## Notes

### Competing Interest Statement

The authors have declared no competing interest.

